# Effect of dioxins in milk on 3D-cultured primary buffalo granulosa cells, a pilot study for a prospective RT-LAMP colour reaction for dioxin toxicity

**DOI:** 10.1101/2020.03.27.011726

**Authors:** Chhama Singh, Mamta Pandey, Emmagouni Sharath Kumar Goud, Vedamurthy G. Veerappa, Dheer Singh, Suneel Kumar Onteru

## Abstract

Dioxins are highly toxic environmental persistent organic pollutants. In several countries, their presence was also reported in cow and human milk samples in the range of 0.023-26.46 and 0.88-19.0 pg/gm of fat, respectively. The detection of dioxns in food samples has been relied on several expensive technologies, which do not represent their toxic effects on consumers. However, mammalian cell based bioassays have such potential to detect the toxins, while representing their toxic effects. Therefore, we tried a three-dimensionally (3D) cultured buffalo granulosa cell based RT-LAMP colour reaction for detecting the presence of added model dioxin, 2,3,7,8-tetrachlorodibenzodioxin (TCDD), in commercial milk. The 3D spheroids on the fifth day of culture were treated with different concentrations of TCDD (i.e. 0.02– 20 pg/ml) directly as well as indirectly through milk fat. After 24hrs of treatment, gene expression studies were performed on certain granulosa cell-specific (*CYP19A1, ER-beta, FSHR* and *LHR*) and selective TCDD-responsive (*CYP1A1, CYP1B1* and *AHR*) genes to identify the potential dioxin responsive gene for further RT-LAMP reaction. As the *AHR* expression in 3D cultured buffalo granulosa cells appears to be a potential gene marker for sensing the added TCDD in the milk, a colour based RT-LAMP reaction was successfully attempted for its expression. However, future studies are needed to develop a dose-responsive colour reaction by considering the treatment time less than 24 hrs.

## 1. Introduction

Dioxins and dioxin-like compounds (DLCs) are considered as one of the highly toxic environmental persistent organic pollutants (POPs). Around 419 types of dioxins and dioxin-like compounds have been identified so far, and 30 of them were found to have significant toxicity. Among them, the 2,3,7,8-tetrachlorodibenzodioxin (TCDD) is the most toxic dioxin with a toxic equivalency factor (TEF) of one (Pohjanvirta et al., 1994). Exposure to dioxins causes skin disorders like choric acne and scar (Mocarelli et al., 1991), nervous disorders, carcinogenesis, generative dysfunction, suppression of the immune system, and imbalance in the sex ratio of the newborn in humans (Ferrari et al., 2000). The predominant human exposure to dioxins is through food, especially fat from meat, fish and dairy products (https://www.who.int/news-room/fact-sheets/detail/dioxins-and-their-effects-on-human-health). Due to lipophilic nature, dioxins get accumulated in the adipose tissue and milk fat. The levels of dioxins in breast milk are higher than in maternal blood (Fukata et. al., 2005). In several countries, dioxin levels were detected in cow and human milk in a range of 0.023-26.46 pg/gm of fat and 0.88-19 pg/gm of fat, respecively. The feeding of ruminants on the contaminated feed materials (Jensen, & Hummel, 1982), including improperly burnt plastic waste which is prevalent in the urban areas of developing countries, results in the engrossment of dioxins in milk

Several expensive, time consuming and less sensitive detection methods are available for the detection of dioxins in food materials. The drawbacks of the currently available dioxin detection methods were well documented elsewhere (Tian, et al., 2012). Among those several dioxin detection methods, HRGC/HRMS (High-Resolution Gas Chromatography/ High-Resolution Mass Spectroscopy) is the most common popular method. The q-PCR (quantitative polymerase chain reaction), EROD (7-ethoxyresorufin-O-demethylase), CALUX (chemical-activated luciferase gene expression), CAFLUX (chemical-activated green fluorescent luciferase gene expression), and GRAB (gel retardation of AhR) are the cell-based bioassays that depend on the AhR signaling pathway, a key pathway for dioxin mechanism of action in cells (Tian *et.al.*, 2012). The other methods are the immune assay methods like ELISA (enzyme-linked immunosorbent assay) and Ah-I (active aryl hydrocarbon immunoassay). However, their use is often prohibited due to their high background noise with high false-negative results. Therefore, there is a need for a rapid and cost-effective colour based dioxin detection method representing dioxin toxicity inappropriate mammalian cell model systems. The present manuscript deals with the possibility of 3D-cultured buffalo granulosa cells as a mammalian cell model system to prepare the potential colour based reaction for the dioxins in the milk.

It is well known that human single copy protein orthologs repertoire showed more similarity to the cow protein orthologs than the common laboratory animals like mice and rats (Elsik *et.al*., 2009). However, cattle tissues cannot be obtained in India due to a ban on their slaughtering. Hence, buffalo cells could be alternative mammalian model systems as the buffalo is the closest species to cattle (Glanzmann et al., 2016). Being distant and present in a more protected environment in ovarian follicles, granulosa cells might be more sensitive to food toxicants. Thus, we hypothesize that they can be used for toxicity testing.

Our hypothesis is that the granulosa cells when exposed to toxicants could alter the expression of several genes including the toxicity representing genes, such as *CYP1A1, CYP1B1, AHR*, and granulosa cell specific genes, such as *ER-β, FSHR, LHR*, and *CYP19A1*. Hence, RNA markers could represent the toxicity of most of the toxicants, including dioxin. However, there is no colour based bioassay available using these RNA markers and granulosa cells for the detection of dioxins in milk. Therefore, the present work was focused on granulosa cells, considering them as the best representative of the reproductive toxicity by dioxins. In our previous study, we had developed CYP19-RT-LAMP assay for mammalian tanscript analysis in low-input laboratories (Pandey, *et al.*, 2018) by using 3D cultured buffalo granulose spheroids treated with lipopolysaccharide (LPS). It is evident by several earlier studies that the LAMP technique was more specific and sensitive without the influence of other non-targeted nucleic acids and PCR inhibitors present in the tested samples, like blood and food materials. This high specificity could be due to its specififcity of primers (Inacio et al., 2008). However, this technique was not yet used for the development of cell-based bioassay for the detection of dioxins in milk. Therefore, the current work attempts to develop a colour based assay for dioxins in milk using RT-LAMP and buffalo granulosa cells.,

## 2. Material and methods

### 2.1. 3D-culture of buffalo granulosa cells

Ovaries were collected from buffaloes immediately after slaughtering at slaughter house, Delhi, India, and they were brought to the laboratory in normal saline containing Streptomycin (250 ng/ml) and Penicillin (100 U/ml) within 4 h at 4 **°**C. The processing of ovaries and the granulosa cells isolation method were already established in our laboratory (Yadav et al., 2017). Briefly, the follicular fluid from small (< 4mm diameter) ovarian follicles was aspirated with 5 ml syringe and 18 gauge needle. The fluid was collected into sterile phosphate buffered saline (PBS) containing 100 U/ml of penicillin, 250 ng/ml of streptomycin, 2.5 µg/ml of amphotericin B (Sigma, USA) and 50 µg/ml of gentamicin (Sigma, USA). The fluid containing GCs were centrifuged at 1400 rpm for 5 min to pellet the cells. The GC cell pellet was further suspended in 1-2 ml of serum free Dulbecco’s Modified Eagle Medium (DMEM) (Sigma, U.S.A) supplemented with 3 mM/ml of L-glutamine (Sigma, Japan), 1 mg/ml of bovine serum albumin (Sigma Aldrich, USA), 4 ng/ml of sodium selenite (Sigma, Germany), 2.5 µg/ml of transferrin (Sigma, USA), 2 µM of androstenedione (Sigma, USA), 10 ng/ml of bovine insulin (Sigma, USA), 1.1 mM of MEM non-essential amino acid 100X (Sigma, USA), 1 ng/ml of ovine FSH (Sigma, USA), 10 ng/ml of human rIGF1 (recombinant Insulin growth factor-1) (Sigma, USA), 10 ng/ml amphotericin B, 100 U/ml of penicillin and 250 ng/ml of streptomycin. Cell counting was performed on haemocytometer counting chamber by using trypan blue dye exclusion method. The isolated granulosa cells were seeded in the wells of polyHEMA (Poly 2-hydroxyethyl methacrylate) (Sigma, USA) coated 6 well plates at a density 6 × 10^5^ viable cells/ml. To prepare polyHEMA solution (1X solution = 12 mg/ml), 0.12 g of polyHEMA was weighed and dissolved in 10 ml of 95% ethanol. A total of 700 µl of polyHEMA solution was poured in 35 mm diameters wells of 6-well plate and kept undisturbed at room temperature until the solution got completely evaporated. The cells at a density of 20 × 10^5^ viable cells/ml were also cultured in 10µl hanging drops on 6-well culture plate and incubated at 37 **°**C temp, 5% CO_2_ and 95% relative humidity. The cultures were maintained for 6 days and the media were changed or supplemented after every 48 h in polyHEMA coated plates and hanging drops, respectively (Supplementary figures 1 and 2).

### 2.2. Milk fat extraction

The fat from cow milk was extracted by Dichloromethane/ethanol extraction method (Stefanov et al., 2010). Cow milk from Mother Dairy brand, India, was taken as a sample and spiked with TCDD (2 ng/40 mg of Fat or 1 ml of milk with 4% fat) and incubated for 1 h. Later, 1.5 ml of Dicholoromethanol and ethanol 2:1 mixture (DM-E) solution was added to 1 ml of TCDD spiked milk as well as unspiked milk (control) samples. The mixture was vortexed for 90 sec and centrifuged at 2500 × g for 8 min at −4 °C. The top aqueous phase was pipetted out and discarded. The remaining solution was once again mixed with 1ml of the DM-E by vortexing. The content were centrifuged at 2500 × g for 6 min at −4°C and obtained three layers. The bottom layer was comprised of milk protein precipitate, the middle layer was of an aqueous phase and the upper layer was of organic phase containing the milk fat. The upper organic phase was collected in another microcetrifuge tube and evaporated to complete drying. The dried fat was then dissolved in 1% DMSO (Dimethyl sulphoxide).

### 2.3. Milk fat measurement

Milk fat was measured by a simple UV spectrophotometric method (Forcato et al., 2005). Briefly, 3 ml of absolute ethanol was added to 60 µL of milk and incubated at −20 °C for 1 h to allow the precipitation of proteins and hydrophobic peptides, which usually interfere in measurement of fat under UV light. After incubation, the mixtures were centrifuged at 13,000 rpm for 15 min. The absorbance of the supernatants were measured in 1-cm path quartz cuvettes under a spectral range of UV wavelengths from 200 to 300 nm. It was noted that the maximum absorbance at 208 nm. The fat percentage was then calculated according to Forcato et al. (2005) by considering the representation of the absorbance value as 1.5 at 208 nm towards 2000 µg of fat per 60 µl of milk.

### 2.4. Treatment of granulosa cell spheroids with TCDD through media and TCDD through milk fat in media

The buffalo granulosa cell spheroids (Supplementary Figures 1 and 2) on the 5^th^ day of the culture were treated with different concentrations of TCDD directly in media or indirectly with the milk fat containing TCDD. Primarlily, the working solution of TCDD was prepared by diluting the TCDD stock solution (10µg/ml of toluene) (CHEMSERVICE.inc;Cat.no: S-10607U10-1ml) into 10ng/ml of 1% DMSO. Later, the working solution was directly added to the media at different concentrations, such as 0.02 pg/ml, 0.2 pg/ml, 2 pg/ml, 10 pg/ml and 20 pg/ml of media, to treat the granulosa cells spheroids. In order to treat the granulosa cell spheroids indirectly with TCDD through milk fat, another stock TCDD solution was prepared by adding 2 ng of TCDD to 1ml of milk containing 4% fat, and the TCDD spiked milk was incubated at room temperature for an hour. With an assumption that all the extracted fat would recover the added TCDD, total milk fat was extracted from TCDD spiked milk using dichloromethane-ethanol extraction method. The extracted fat was dissolved in 1 ml of 1% DMSO, which was used to treat the granulosa cell spheroids at different concentrations, such as 0.02 pg/ml, 0.2 pg/ml, 2 pg/ml, 10 pg/ml and 20 pg/ml of media (Goud, *et al.*, 2019).

### 2.5. RNA isolation and cDNA synthesis from cultured granulosa cell spheroids

Total RNA was isolated from the cultured granulosa cell spheroids by using Trizol reagent (Sigma-Aldrich) according to the manufacturers instructions. The isolated RNA was either immediately used for reverse transcription polymerase chain reaction (RT-PCR) or stored at −80 **°**C for further use. The RNA was quantified on Nano Quant (IMPLEN Photometer) and the cDNA was synthesised from 200ng of total RNA by First-Strand cDNA synthesis kit (CAT#k1622). The prepared cDNA was diluted 10 times prior to its usage in qRT-PCR.

### 2.6. Relative quantification of the selected dioxin responsive and GC specific transcripts

Quantitative real time PCR (qRT-PCR) was used to quantify the expression of the selected dioxin responsive and GC specific transcripts by considering the ribosomal protein lateral stalk subunit P0 (*RPLP0*) transcript as a housekeeping gene. The primers specific to the selected genes were designed using Primer Blast Software (http://www.ncbi.nlm.nih.gov/tools/primer-blast) (Supplementary Table 1). The RT-PCR reaction was performed in 12 µl reaction mixture, which was prepared by mixing 5 µl of SsoFast™Evagreen® Super Mix, 5 µl of cDNA, 1µl of forward primer (5 µM) and 1µl of reverse primer (5 µM). The qRT-PCR conditions were as follows: initial denaturation at 95°C for 5 min, followed by 40 cycles of denaturation, annealing and extension at 95°C for 10 sec, 60°C for 30 sec and 72°C for 30 sec, respectively. To further ensure the amplification of the PCR products, the meltcurve peaks were obtained finally by exposing the PCR products to 95°C for 5 sec, 60°C for 1 min, 97°C for 1 min and final cooling at 40°C for 30 sec.

**Table 1.**
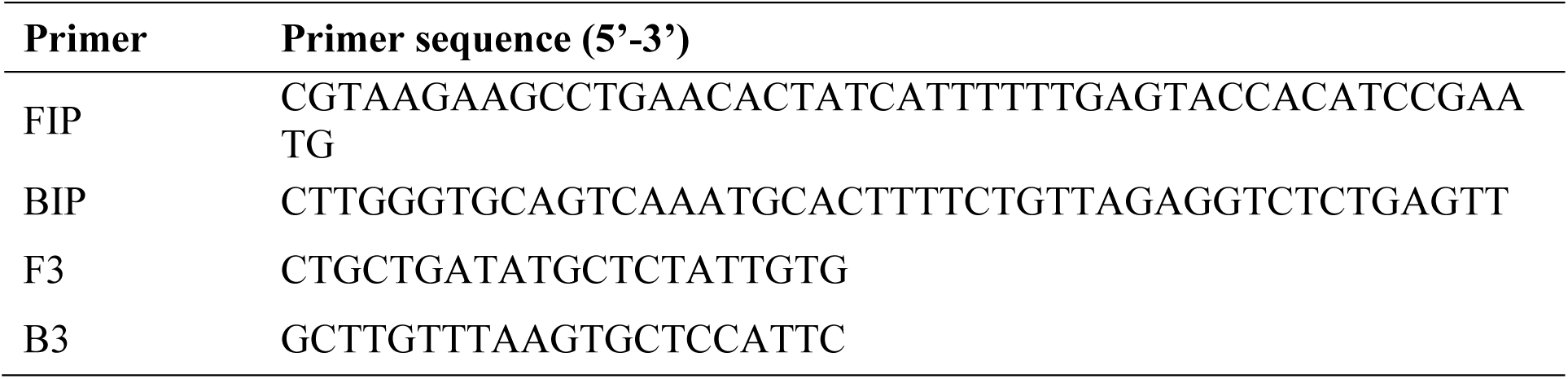
LAMP primers for *AHR* gene.

### 2.7. Statistical analysis

Statistical differences among the mean foldchages of each gene expression at different treatment groups were identified by one-way ANOVA with posthoc Tukey test using Graph Pad Prism software version 5.01 (Graph Pad Software Inc., San Diego, California, USA). The P < 0.05 was considered as statistically a significant difference among the treatment groups. Furthermore, a heat map was generated using the HEATMAPPER (http://heatmapper.ca/) for the mean fold changes of all the selected transcripts at different treatments groups in two different culture systems. Additionally, correlation values were computed for each gene using its fold change values between the direct TCDD treatment groups and the groups treated with TCDD through milk fat under a culture system. The computed correlation values were used to select the best TCDD responsive transcript, whose expression dynamics was not affected by the milk fat.

### 2.8. Reverse-transcription Loop Mediated Isothermal Amplification (RT-LAMP)

The above correlation analysis identified that AHR gene could be the best TCDD responsive transcript. Hence, LAMP primers such as forward inner (FIP), backward inner (BIP), forward outer (F3) and backward outer (B3) primers (Table 1) were designed for the AHR gene using PrimerExplore V5 software. The RT-LAMP reaction was performed in a 25 µl PCR system comprising 12.5 µl of Warmstart 2X Master Mix, 1.6 µM each of FIP and BIP, 0.2 µM each of F3 and B3, 120 µM of HNB, 6 mM of MgSO_4_ and the specific amount of RNA template. The reaction conditions were initial incubation at 65°C for 1 hr and later at 80°C for 10 min to terminate the reaction.

## 3. Results and disscusion

### 3.1. Milk fat estimation

The fat percentage for all the milk samples was found to be nearly 4.4 %. Precisely, the fat percentage of liquid milk, extracted milk fat, and extracted milk fat from TCDD spiked milk samples was 4.24±0.144 gm/100 ml, 4.472±0.248 gm/100 ml and 4.4075±0.249 gm/100 ml, respectively.

### 3.2. Relative quantification of the selected TCDD responsive genes

In the present study, the expression of three TCDD responsive genes (*CYP1A1, CYP1B1* and *AHR*) was studied. The *CYP1A1* expression was significantly (P < 0.05) higher in buffalo granulosa cell spheroids treated with the milk fat isolated from the spiked milk containing 0.2 pg/ml of TCCD than those treated with milk fat without TCDD (Figure 1 and Supplementary figure 3) in both the 3D culture systems. In rat granulosa cells, the *CYP1A1* gene expression was detectable only after frequent treatments with higher doses of TCDD (3.1nM) during 6 hr of the culture, but not at 24 hr or 48 hr of culture (Dasmahapatra et al., 2001). In the present study, a long duration (24 hr instead of 6 hr) treatment with 0.2 pg/ml of TCDD through milk fat may be the reason for significantly higher expression of the *CYP1A1*. Although there was an increasing trend of *CYP1B1* expression by the milk fat isolated from the milk spiked with TCDD, there was no significant difference between milk fat and milk fat containing TCDD treatments in both the culture systems (Figure 1 and Supplementary figure 4). This could be due to adaptive tolerance of buffalo granulosa cell spheroids towards milk fat TCDD treatment for 24 hr. Taken together, the expression of the *CYP1A1* and *CYP1B1* indicated both the time and dose dependent manner for TCDD treatment through milk fat, as TCDD greater than 0.2 pg/ml in milk fat might reduce their expression levels in buffalo granulosa cell spheroids during 24 hr treatment.

**Figure 1.**
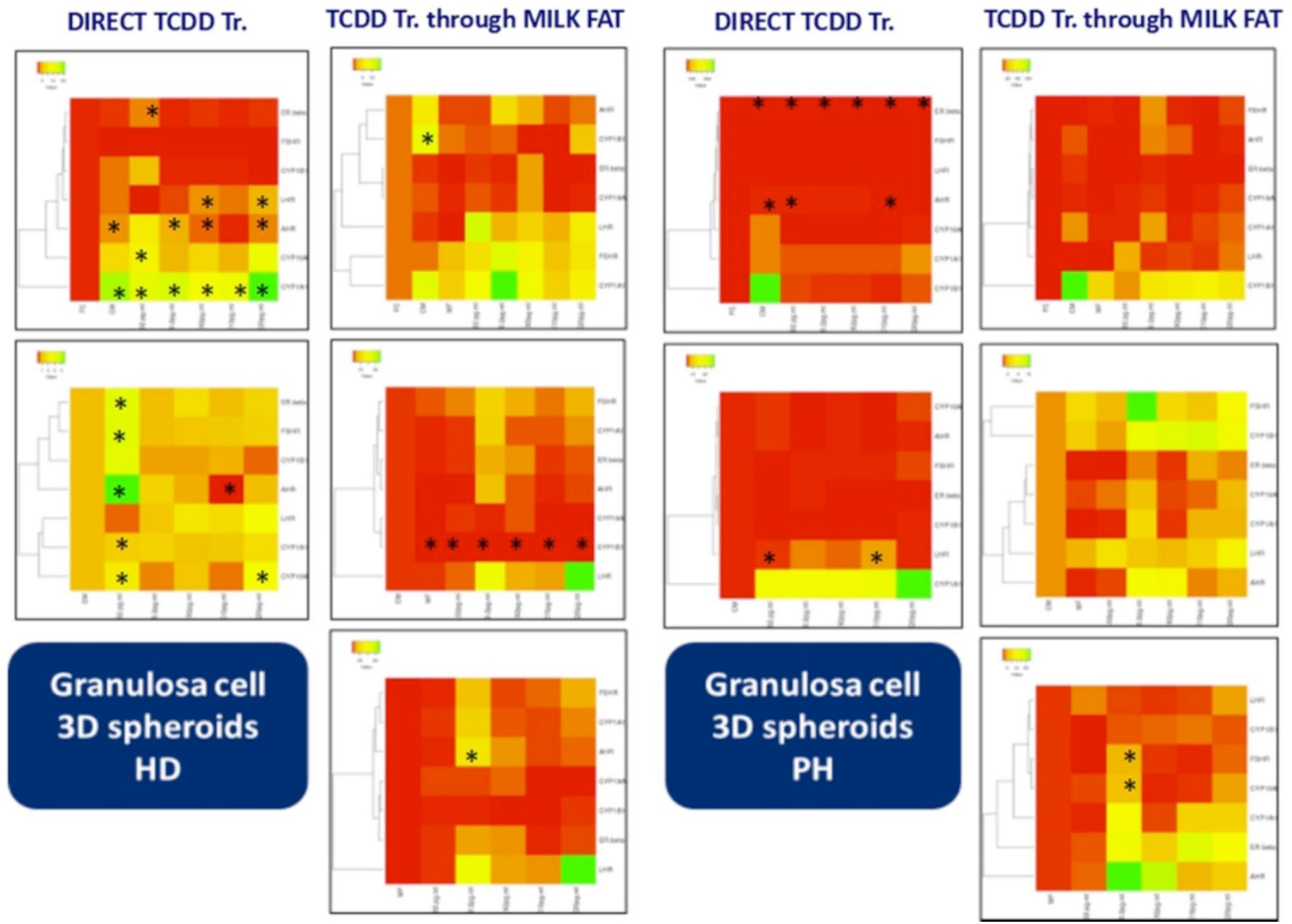
Heatmap for the gene expression of the selected TCDD responsive genes and candidate granulosa cell markers. The average fold changes of the gene expression of the selected candidate granulosa cell markers i.e., *CYP19A1, ER-β, FSHR* and *LHR*, and the selected TCDD responsive genes i.e., *CYP1A1, CYP1B1* and *AHR*, in primary buffalo granulosa cells spheroids cultured in hanging drop and polyHEMA culture system were depicted in a heatmap by the heaptmapper software. The No.s 1 and 2 represent the experiment results from hanging drop (HD) culture system. The No.s 3 and 4 represent the experiment results from polyHEMA (PH) culture system. The No.s 1 and 3 represent the treatment of spheroids with direct TCDD treatment. The No.s 2 and 4 represent the treatment of spheroids with fat isolated from TCDD spiked milk. CM represents the cells in media. 0.02pg/ml, 0.2pg/ml, 2pg/ml, 10pg/ml and 20pg/ml indicate the conc. of TCDD. FC represents granulosa cells isolated freshly. MF represents milk fat. The colour pattern from red to green indicates increased gene expression and the * indicates the significant difference in the gene expression among different concentrations of TCDD treatments.

The increased expression of the *CYP1A1* and *CYP1B1* genes in response to TCDD treatment indicated the activation of ovarian *AHR* signalling pathways. The *AHR* gene expression was significantly higher in buffalo granulosa cell spheroids treated with the milk fat containing 0.2 pg/ml of TCDD than the milk fat alone in hanging drop 3D culture system. A similar trend of the higher expression of *AHR* gene, although non-significant, was observed by the milk fat containing either 0.2 or 2 pg/ml of TCDD in the polyHEMA culture system (Figure 1 and Figure 2). These observations indicated that buffalo granulosa cells could be used to detect dioxins in the milk through *AHR* gene expression. However, the treatment period (24 hr) of the granulosa cell spheroids need to be decreased to obtain a dose dependent standard curve. By 24 hr treatment with higher concentration of TCDD (> 2pg/ml of TCDD in milk fat), the cells might have developed tolerance or non-responsiveness, which limit their use for biosensor development.

**Figure 2.**
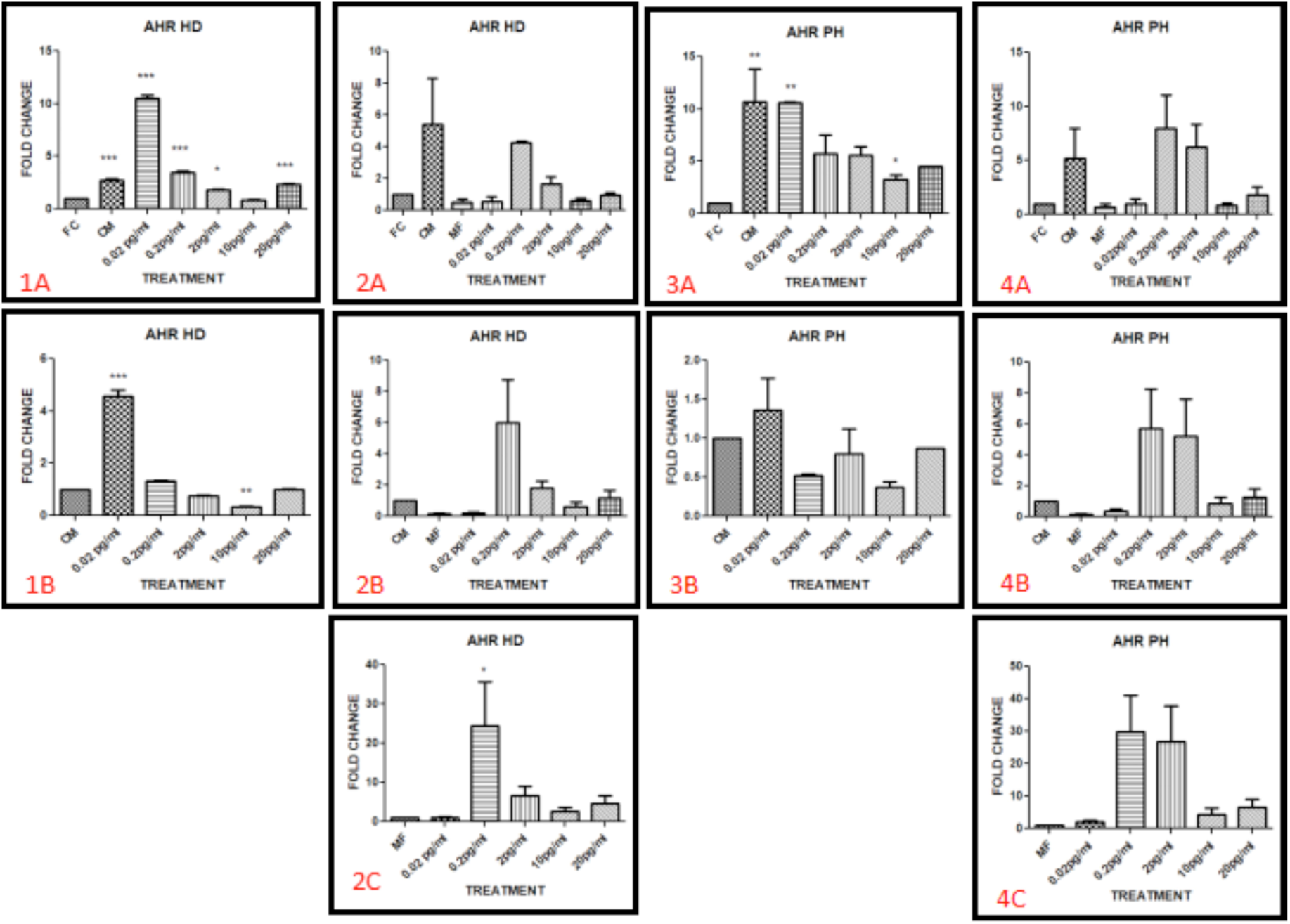
Fold change graph of *AHR* gene in Hanging drop and polyHEMA culture system for primary buffalo granulosa cells spheroids 3D culture. When the buffalo granulosa cell spheroids in hanging drop culture system were treated with TCDD directly in the media, the expression of the *AHR* gene was significantly higher at 0.02pg/ml TCDD concentration than fresh and control cultured spheroids, and then the expression declined at other concentrations of TCDD treatment. A similar increasing trend of *AHR* gene expression at 0.02pg/ml of TCDD concentration and a decreasing trend of its expression at other concentration of TCDD directly present in the media were observed in polyHEMA culture system. When the spheroids were treated with the milk fat containing TCDD, the *AHR* gene expression was significantly high when the milk fat contained 0.2pg/ml of TCDD in hanging drop system. A similar trend of higher expression of *AHR* gene, although non-significant, was observed at milk fat treatment containing either 0.2 or 2 pg/ml of TCDD in the polyHEMA culture system. 1 and 2 represents experiment results in hanging drop (HD) culture system.3 and 4 represents experiment results in polyHEMA (PH) culture system.1 and 3 represent treatment of spheroids with TCDD. 2 and 4 represents treatment of spheroids with fat isolated from TCDD spiked milk. CM represents the cells in media. 0.02pg/ml, 0.2pg/ml, 2pg/ml, 10pg/ml and 20pg/ml indicate the conc. of TCDD. FC represents granulosa cells isolated freshly. MF represents milk fat.

Significantly increased *AHR* gene expression was also observed in 2D cultured rat granulosa cells by direct TCDD-treatment for 48 hr (Dasmahapatra et al., 2001). Generally, the *CYP1A1* gene expression was considered as a biomarker for the activation of the *AHR* by TCDD. However, time-dependent expression of the *AHR* and *CYP1A1* genes was observed in mouse ovarian cancer cell line ID8 in response to TCDD treatment. The increased *AHR* gene expression coincides with the maximum induction of *CYP1A1* gene till 6 hr of TCDD treatment, but not in the later period of the treatment (Son et al., 2002). In the current study, TCDD treatment in the milk fat increased the *AHR* expression upto 0.2pg/ml and then decreased at 20 pg/ml in 24 hours (Figure 1 and Figure 2). These observations indicated that the *AHR* expression in granulosa cells could be species, dose and time dependent manner with TCDD treatments. This kind of different AHR gene expression under different TCDD treatments could be due to the differences between the AHR mRNA and protein levels. In porcine granulosa cell culture, TCDD decreased *AHR* protein expression after 1hr and 48 hr, but AHR mRNA expression was influenced only after 48 h (Piasecka-Srader et al. 2016). This shows that TCDD induces rapid reduction of AHR protein prior to its significant effect on AHR mRNA expression in *in vitro* cell culture systems. It further strengthens that there may be a variation in TCDD’s influence on the gene expression in different tissues, species and experimental conditions.

### 3.3. Relative quantification of the granulosa cells specific markers

The expression of ER-beta gene was increased in a group treated with 0.02 pg/ml of TCDD directly in hanging drop culture system or greater than 0.2 pg/ml of TCDD in milk fat in both the culture systems (Supplementary figure 5). This expression is akin to *AHR* expression by the TCDD in milk fat (Figure 2). These observations indicate that the *ER-beta* could be another potential gene marker like *AHR* gene to identify the dioxins in milk. This similarity in the *ER-beta* and *AHR* expression could be due a crosstalk between AHR and estrogen receptors, particularly ER-beta rather than ER-alpha (Piasecka-Srader et al., 2016). It is well known that the granulosa cells of small follicles in most species including rat and human express *ER-beta* as a dominant isoform of estrogen receptors in their nuclei (Chu et al., 2004). In the present study also, the granulosa cells from small ovarian follicles of buffaloes were used. The increased expression of ER-beta in buffalo granulosa cells by the TCDD treatment is similar to the earlier studies in rat granulosa cells (Dasmahapatra et al., 2000). TCDD induced *ER-beta* and *AHR* mRNAs in cultured rat granulosa cells at 48h, and this induction might be due to the catabolism of estrogens by the *CYP1A1*/*CYP1B1* gene products through the activation of the AHR gene. In fact, ER-beta mRNA levels were increased at the plateau stage of the CYP1B1 mRNA expression by the nM of TCDD treatment to rat granulosa cells (Dasmahapatra et al., 2001). In contrast, *in-vivo* studies in mice have shown significantly decreased *ER-beta* mRNA in the ovary (*p*<0.05) and uterus (*p*<0.01) with TCDD treatment of dose 5 *µ*g/kg body weight (Tian et al., 1998). Similarly, constitutively activated *AHR* gene showed an inhibitory effect on estrogen synthesis in KGN cells, a human granulosa cell line (Horling et al., 2010). All these observations seem that TCDD’s influence on *ER-beta* expression varies with different doses, cell types and species.

In the current study, the *CYP19A1* gene expression was increased in the cultured granulosa cells in hanging drop and polyHEMA culture system after 24 hr exposure with 2 pg/ml and 0.2 pg/ml of TCDD in milk fat (Supplementary figure 6). On the contrary, TCDD significantly reduced the *CYP19A1* gene expression, but not its protein levels, in human choriocarcinoma cell line, JEG-3, at 48 h and 96 h (Drenth et al., 1998, Karman et al., 2012). In rat granulosa cells, TCDD (3.1nM) did not reduce the aromatase activity at 6 h and 24 h, but reduced at 48 h (Dasmahapatra et al., 2001). However, aromatase enzyme activity was significantly increased by FSH in the culture of prepubertal and adult rat granulosa cells than the control or TCDD at 24 and 48 h. The increased aromatase activity by FSH (50 ng/ml) was significantly inhibited by 3.1 nM of TCDD, but not by the pM doses of TCDD (Dasmahapatra et al., 2000) at 48 h. In the current study, 0.02-20 pg/ml of TCDD was used. This indicated that the inhibited aromatase activity by TCDD could be a dose-dependent process regulated either at the transcriptional level or at the level of post-translational modifications.

In this study, TCDD treatment through milk fat (0.2 pg/ml) showed an increasing trend in the FSHR expression than the treatment by the milk fat alone and the milk fat with other increased concentrations of TCDD in both the hanging drop and polyHEMA culture systems (Supplementary figure S7). Minegishi et al. (1995) reported that FSH treated rat granulosa cell culture resulted in the decrease of *FSHR* level in 2 to 6 hr after the treatment, with a late recovery at 24 hr. On the contrary, simultaneous treatment with FSH and increased doses of TCDD inhibited the *FSH*-induced *FSHR* levels in a dose-dependent manner in rat granulosa cells being cultured from 24 hr to 72 hr. On the contrary, the expression of *LHR* gene was found to be upregulated in a dose dependent manner (not significant) by TCDD in cultured buffalo granulosa cells in hanging drop and polyHEMA culture system (Supplementary figure S8). This is because of the shifting of the gonadotrophin receptor expression towards *LHR* dominance (Horling et al., 2010).

### 3.4. Correlation between TCDD treatment in media and TCDD treatment through milk fat in polyHEMAand hanging drop culture systems

A correlation plot was drawn between the fold changes of direct TCDD treated samples and the samples treated with TCDD through milk fat to choose the best dioxin responsive candidate gene for the preparation of biosensor in both culture systems. Among all the 3 selected TCDD responsive genes (*CYP1A1, CYP1B1* and *AHR*) (Figure 3) and 4 selected granulosa marker genes (*CYP19A1, FSHR, LHR*, and *ER-beta*) (Supplementary figure 9), the *AHR* gene expression showed high positive correlation between the fold changes of TCDD through media treated samples and TCDD through milk fat treated samples at 24 hr in hanging drop culture system. Also, *AHR* gene shows high variation in correlation between 0.02 pg/ml and 0.2 pg/ml. Therefore, we selected *AHR* gene in hanging drop culture system as the best candidate gene to proceed further for RT-LAMP colour assay.

**Figure 3.**
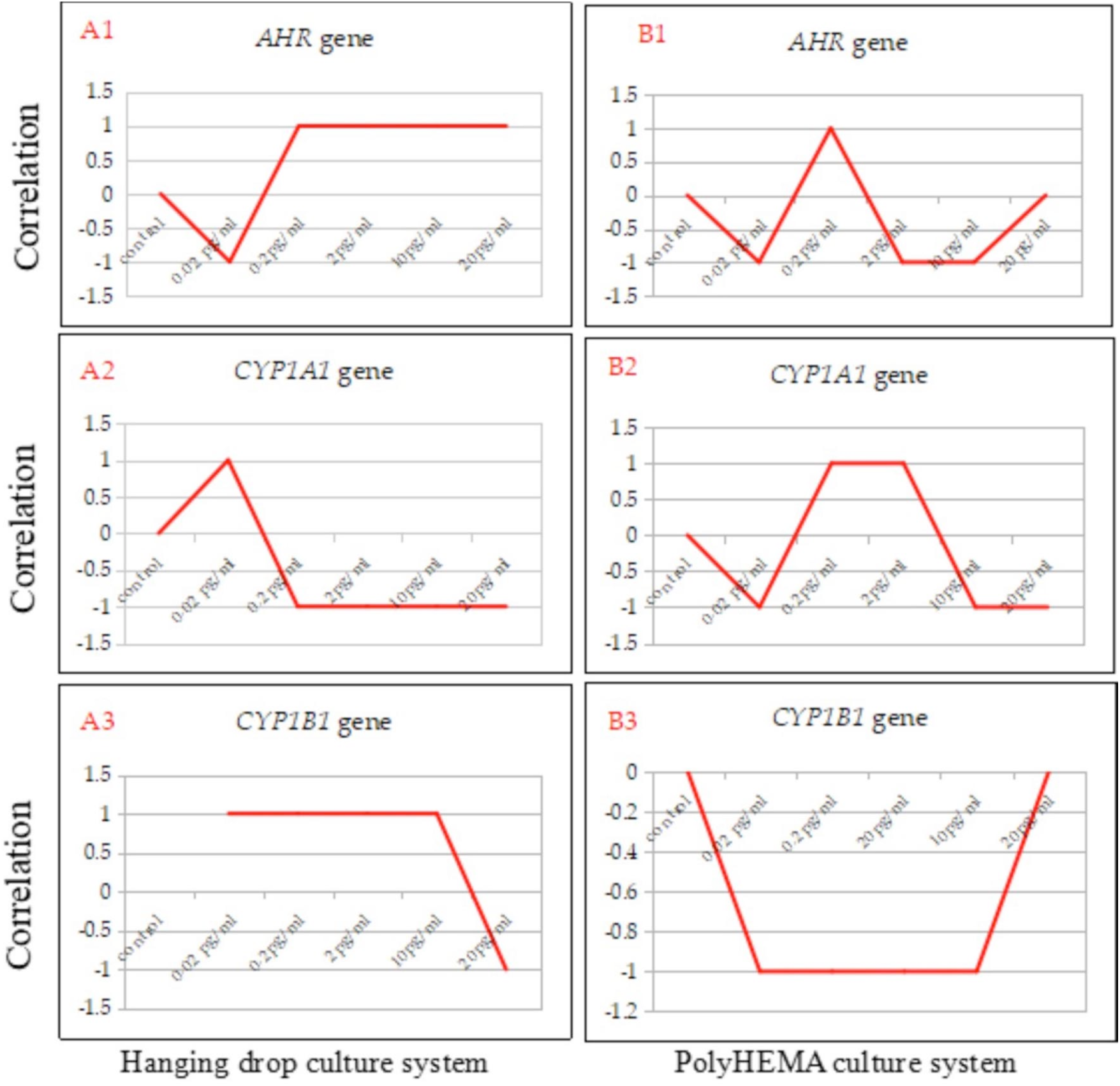
Correlation analysis for the expression of the selected TCDD responsive genes’ between direct TCDD treatment and TCDD treatment through milk fat on 3D cultured primary buffalo granulosa cells spheroids. A and B represent the correlation analysis between the above two kinds of treatments in hanging drop (HD) and polyHEMA culture systems, respectively. The No.s 1, 2 and 3 indicate the *AHR, CYP1A1* and *CYP1B1* gene expressions, respectively. The X-axis shows different concentrations of TCDD and the Y-axis indicates the correlation value between the two kinds of TCDD treatments. Among the selected genes, the *AHR* gene expression has the highest correlation between the two types of TCDD treatments in hanging drop culture system.

### 3.5. Development of RT-LAMP colour reaction for the best dioxin responsive candidate gene

The present study is the first study to use *AHR* gene expression for milk dioxin detection through LAMP reaction, as the amplification products can be detected visually with the naked eye in normal white light. The RT-LAMP was standardized for the amplification of the *AHR* gene from granulosa cells spheroids by standardizing the concentration of template and primers at different temperatures and incubation time according to Pandey et al. (2018). On the basis of colour detection with 120 nM Hydroxynaphthol blue (HNB) dye, all the RT-LAMP positive samples of the *AHR* gene from the granulosa cell spheroids of both the hanging drop and polyHEMA culture systems appeared blue in colour, whereas the non-amplified and non-template controls remained in violet colour (Figure 4 and Figure 5). The hydroxynaphtol blue (HNB) dye, a metal ion indicator, was used in the LAMP assay for visualizing the amplification of the *AHR* gene, as it can be added to the reaction tube prior to the reaction (Balne et al., 2015). In general, different nucleotide intercalating dyes such as SYBR green, picogreen and propidium iodide have been used for better visibility of the LAMP reaction result. But these dyes are usually added to the LAMP reaction after the amplification was completed. Opening the reaction tube to add these dyes may cause contamination from the surroundings to amplified products. To overcome this problem, HNB dye has been successfully used to visualise the LAMP reaction result for *AHR* gene in this study.

**Figure 4.**
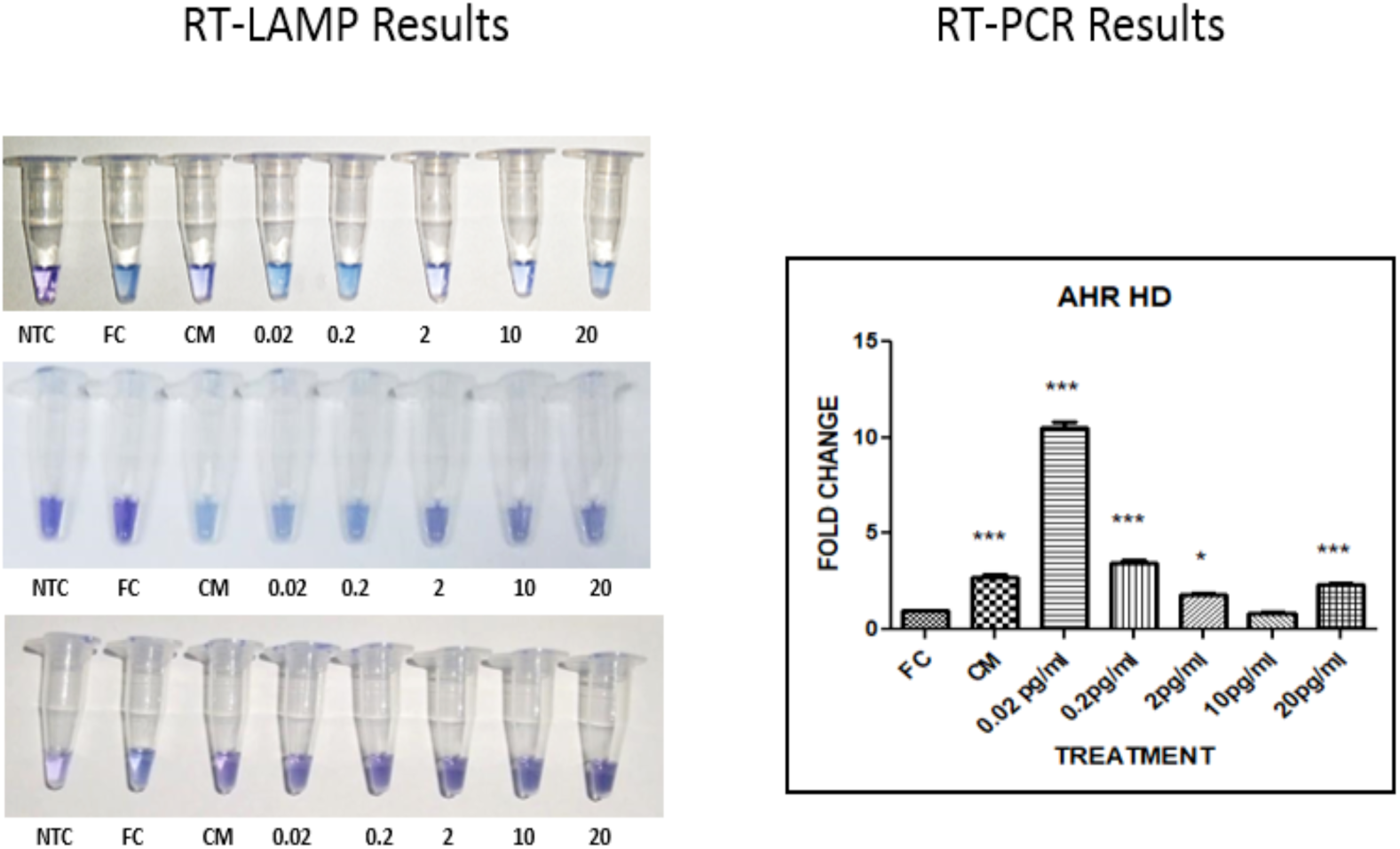
Comparison between RT-LAMP and RT-PCR results for *AHR* gene amplification in 3D cultured primary buffalo granulosa cell spheroids under direct TCDD treatments in hanging drop culture system. NTC, FC and CM indicate non-template control, fresh granulosa cells and 6 days cultured cells without TCDD treatment, respectively. The TCDD concentrations are indicated from 0.02 pg/ml to 20 pg/ml. The colour pattern from violet to blue indicates the increased amplification of the target gene in RT-LAMP products. There was a slight variation between the RT-PCR results and RT-LAMP colour change. The results of RT-PCR were cumulative of triplicates in one figure and RT-LAMP results were depicted as individual single experiment results in triplicates.

**Figure 5.**
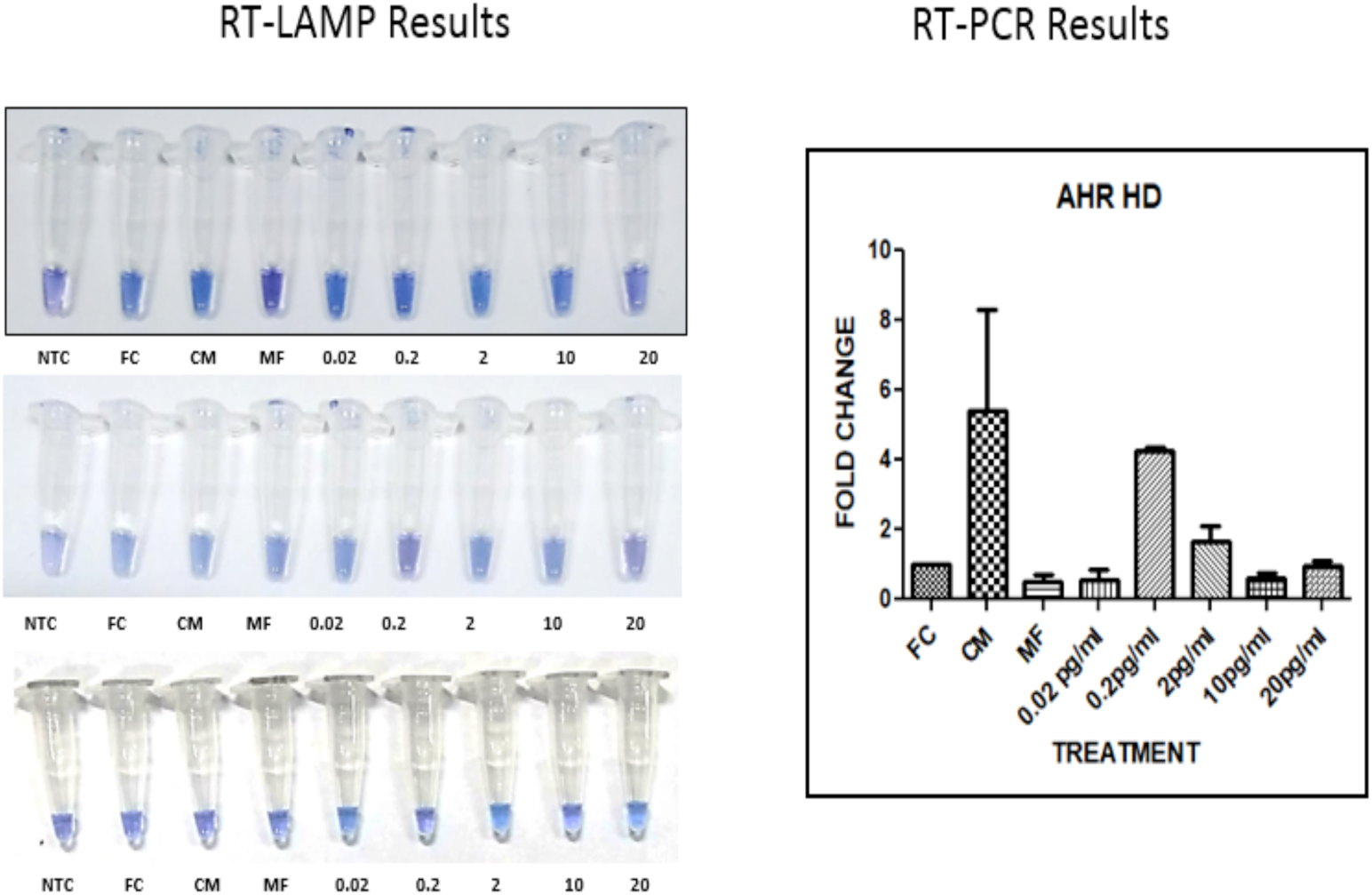
Comparison between RT-LAMP and RT-PCR results for AHR gene amplification in 3D cultured primary buffalo granulosa cell spheroids under treatment TCDD through milk fat in hanging drop culture system. NTC, FC and CM indicate non-template control, fresh granulosa cells and 6 days cultured cells without TCDD treatment, respectively. The TCDD concentrations are indicated from 0.02 pg/ml to 20 pg/ml. The colour pattern from violet to blue indicates the increased amplification of the target gene in RT-LAMP products. There was a slight variation between the RT-PCR results and RT-LAMP colour change. The results of RT-PCR were cumulative of triplicate results in one figure and RT-LAMP results were depicted as individual single experiment results in triplicates. Hence, RT-LAMP needs to be standardized further.

The colour change in LAMP products more or less followed the pattern observed in RT-PCR. For example, high intensity of blue colour showing the increased expression of the *AHR* gene in the LAMP reaction followed the increased expression of the *AHR* gene in the RT-PCR results at 0.02 pg/ml and 0.2 pg/ml of dioxin treatment to granulosa cell spheroids in the hanging drop culture system. There was a slight variation between the RT-PCR results and RT-LAMP colour change, because the results of RT-PCR were cumulative results of triplicate results in one figure and RT-LAMP results were depicted as individual single experiment results in triplicates. That variation was justified by the LAMP reaction for the *AHR* gene expression under TCDD treatment indirectly through milk fat on polyHEMA culture system (Figure 6). The colour of non-template control was violet showing no amplification, and the LAMP product containing RNA of fresh cells as a template showing colour change into blue shows high amplification. The high colour change was observed at 0.02 pg/ml to 0.2 pg/ml of the TCDD treatment. Later at the treatment with higher concentration of TCDD greater than 0.2 pg/ml, a shift of the colour change slightly towards the violet colour indicated lower amplification of the *AHR* gene. Further standardization is required in the future studies for the LAMP reaction towards the granulosa cell based biosensor preparation for dioxins in the milk by considering treatment time less than 24 hrs.

**Figure 6.**
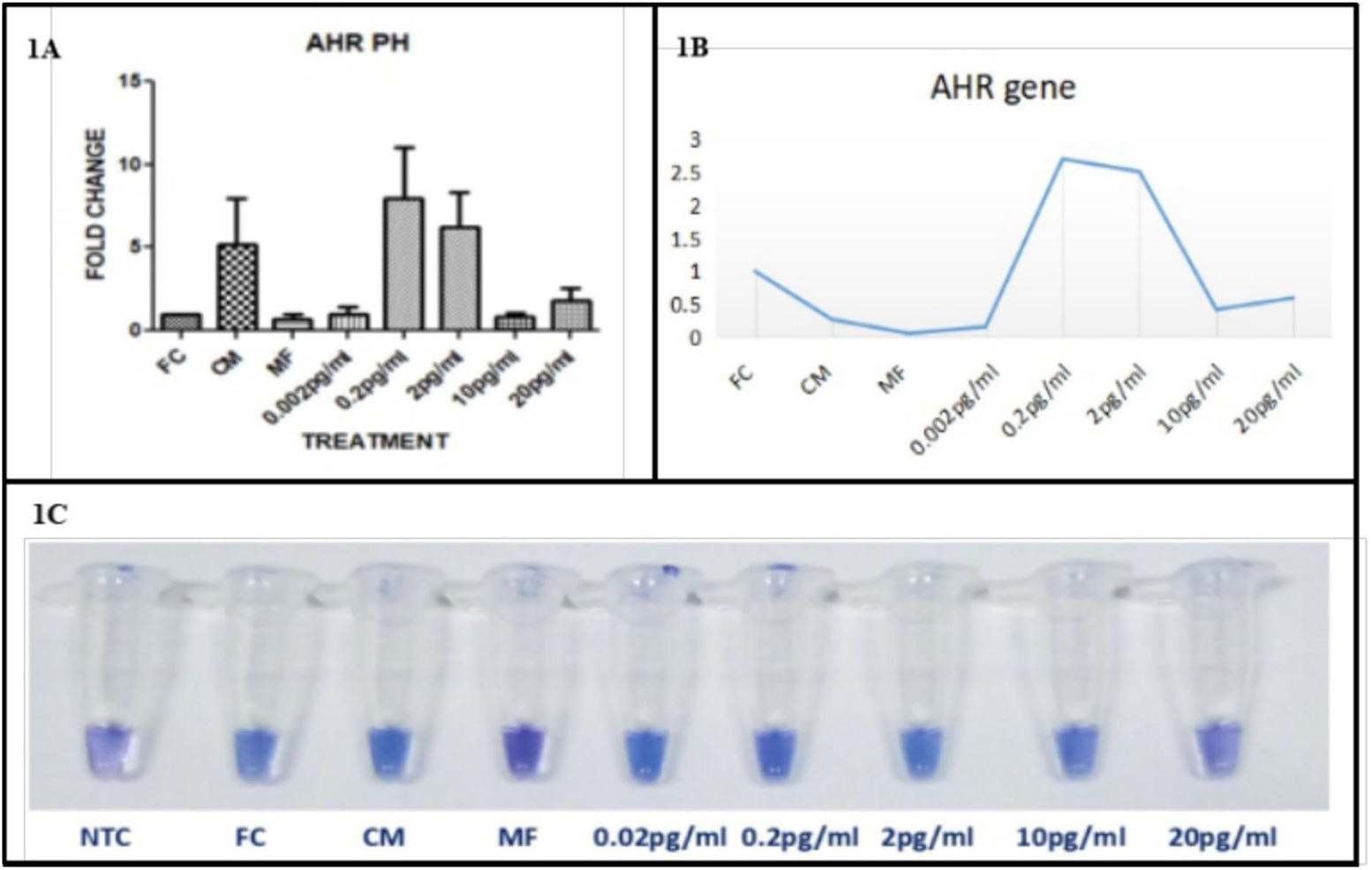
LAMP Reaction for AHR gene expression in buffalo granulosa cells under TCDD treatment through milk fat in polyHEMA systems. NTC, FC and CM indicate non-template control, fresh granulosa cells and 6 days cultured cells without TCDD treatment, respectively. The TCDD concentrations are indicated from 0.02 pg/ml to 20 pg/ml. The colour pattern from violet to blue indicates the increased amplification of the target gene in RT-LAMP products. The variation between the RT-PCR results and RT-LAMP colour change was justified by the LAMP reaction for the *AHR* gene expression under TCDD treatment indirectly through milk fat on polyHEMA culture system. The colour of non-template control was violet showing no amplification, and the LAMP product containing RNA of fresh cells as a template showing colour change into blue shows high amplification. The high colour change was observed at 0.02pg/ml to 0.2pg/ml of the TCDD treatment. Later at the treatment with higher concentration of TCDD more than 0.2pg/ml, a shift of the colour change slightly towards the violet colour indicated lower amplification of the AHR gene.

## 4. Conclusions

The present study is the first attempt to develop a LAMP based reaction for milk dioxins using 3D cultured mammalian reproductive cells, like buffalo granulosa cells. Among the selected TCDD and granulsoa cell specifc genes, the *AHR* gene expression appears to be a potential marker in buffalo granulosa cells for sensing the TCDD in milk. A colour based RT-LAMP reaction was attempted for its expression. However, a dose responsive expression of *AHR* gene needs to be standardized between 0.02 pg/ml to 0.2 pg/ml of TCDD in the milk in future studies.

## Acknowlegement

We thank Director, ICAR-NDRI for providing the infrastructure to carry out the present study. We are highly thankful to the Department of Biotechnology, Ministry of Science and Technology, India, for financial assistance (Grant No. 102/IFD/SAN/3670/2014–15) to this work.

