## Supplementary material for "Effect of dioxins in milk on 3D-cultured primary buffalo granulosa cells, a pilot study for a prospective RT-LAMP colour reaction for dioxin toxicity": supplementary files-2_clean.docx

**SUPPLEMENTARY INFORMATION**

**Supplementary Table 1 - Primer sequences used for real time PCR**

| **S.No.** | **Gene name** | **Primer name: sequence (5’-3’)**  **Forward primer**  **Reverse primer** | **Product length** | **Accession no.** |
| --- | --- | --- | --- | --- |
| 1. | AHR | ATGCTTTGGTTTTCTATGCG  GAGAGTCCATTAGCTTCATCA | 196 | NM_001206026.1  (Bos taurus) |
| 2. | CYP1A1 | CAGGGCGATGATTTCAAGGG  GTCTGAGGCAGTGGAGAAACT | 156 | AB_060696.1  (Bos taurus) |
| 3. | CYP1B1 | CGTTCTCTTCACCAGGTATTC  GTCAAAGTCCTCTGGGTTC | 295 | NM_001192294.1  (Bos taurus) |
| 4. | CYP19A1 | ACTTATCACAACCAGGACTT  GGATGTGTCCTCATAATTCCA | 188 | NM_174305.1  (Bos taurus) |
| 5. | ER-β | TCTCTCCTTTAGCCATCCAT  CTTTTCAATGTCTCCCTGTTC | 121 | XM_006072164.1  (Bubalus bubalis) |
| 6. | FSHR | AGTTGCCCTCTTTCCCATCT  CGCTTGGCTATCTTGGTGTC | 222 | EF650048.1  (Bubalus bubalis) |
| 7. | LHR | GTGTTGCCTCTTGTGGGTGT  GCAGGTGAAATCGGTCAAG | 249 | DQ_858172.1  (Bubalus bubalis) |
| 8. | RPLPO | ATCGCATCTGTACCCCATTC CTCCTTTGCTTCAACCTTGG | 210 | XM_006052971.2  (Bubalus bubalis) |

**Supplementary Figures**

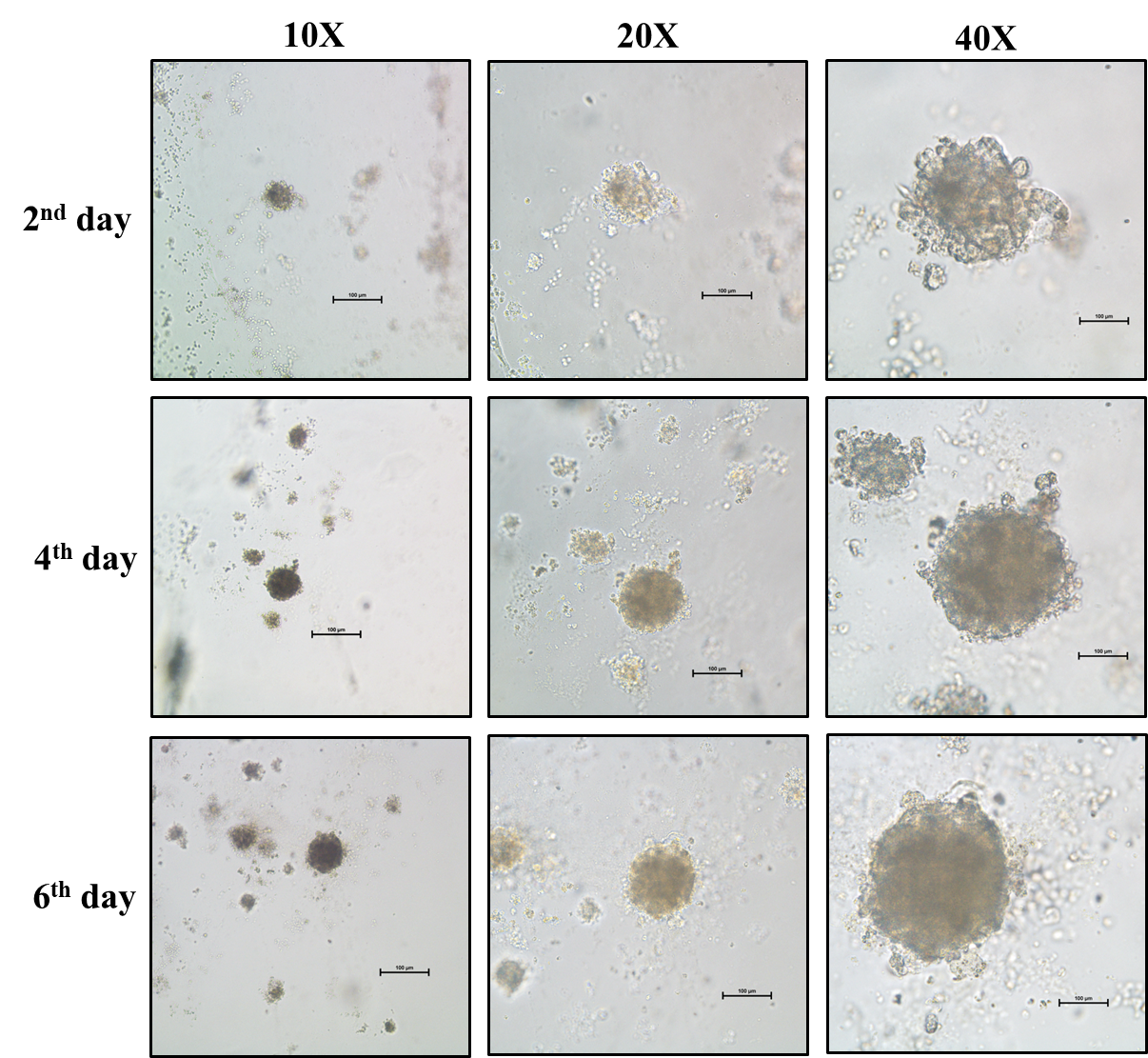

**Supplementary Figure 1.** **Primary buffalo granulosa cells 3D spheroids in hanging drop culture system.** The spheroids became more globular from the 2^nd^ day to 6^th^ day of the culture. The spheroids of buffalo primary granulosa cells on the 2^nd^ day, 4^th^ day and the 6^th^ day are depicted under 10X, 20X and 40X magnifications.

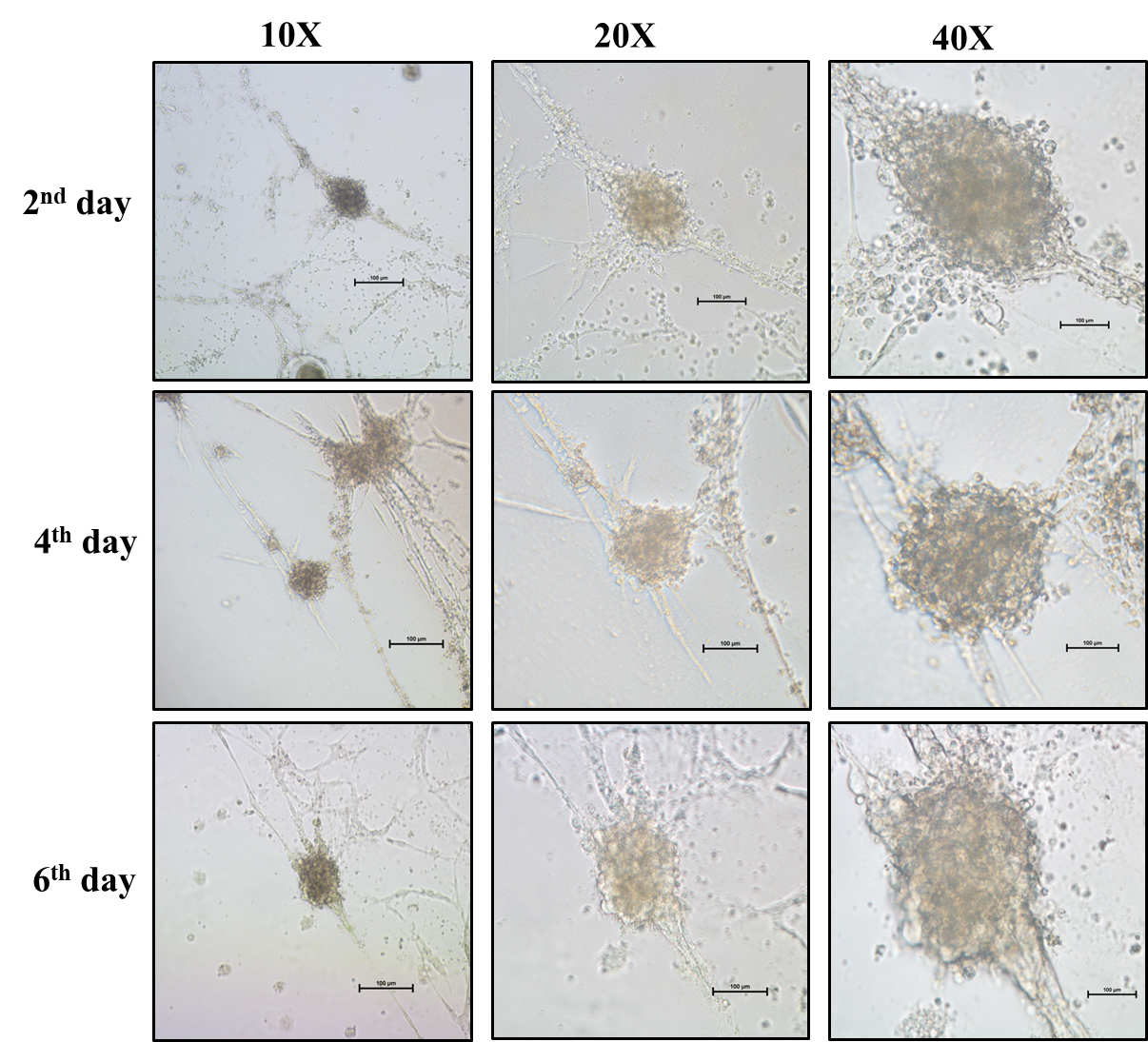

**Supplementary Figure 2.** **Primary buffalo granulosa cells 3D spheroids in polyHEMA culture system.** The spheroids became more globular from the 2^nd^ day to 6^th^ day of the culture. The spheroids of buffalo primary granulosa cells on the 2^nd^ day, 4^th^ day and the 6^th^ day are depicted under 10X, 20X and 40X magnifications.

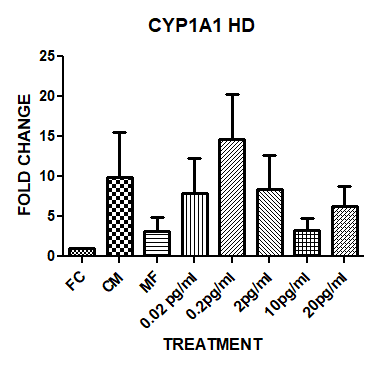

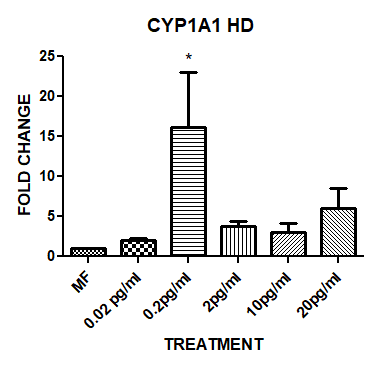

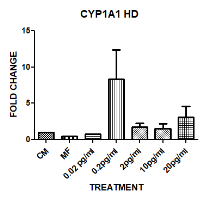

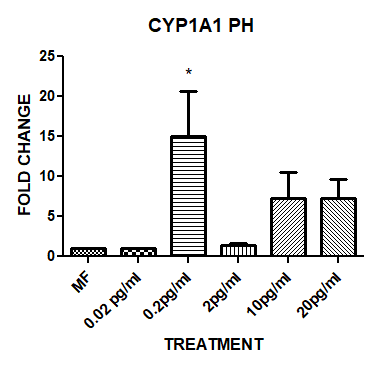

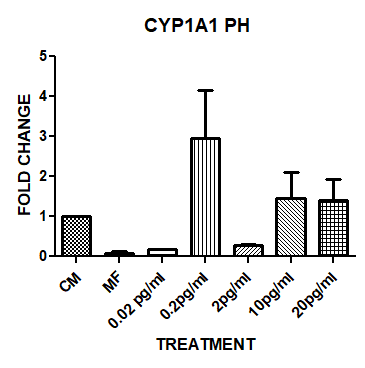

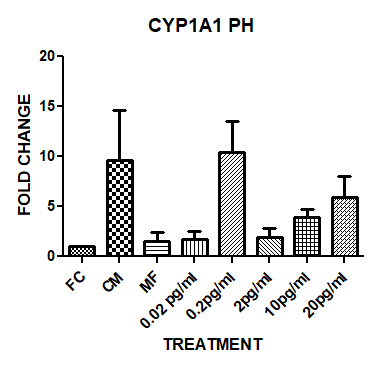

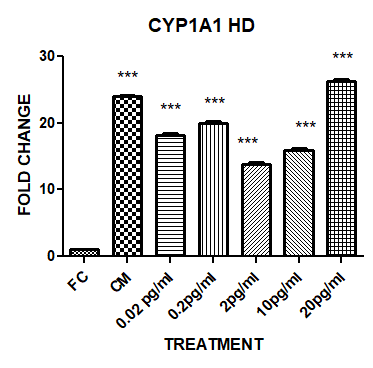

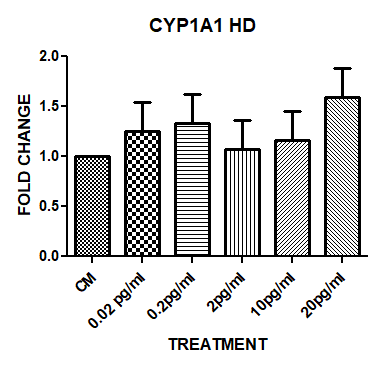

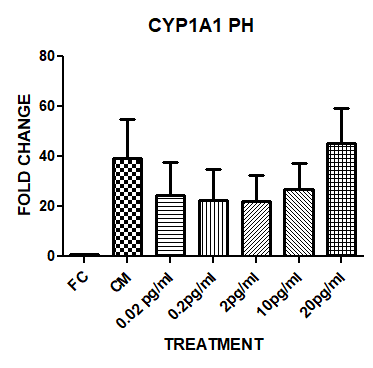

1A

1B

2A

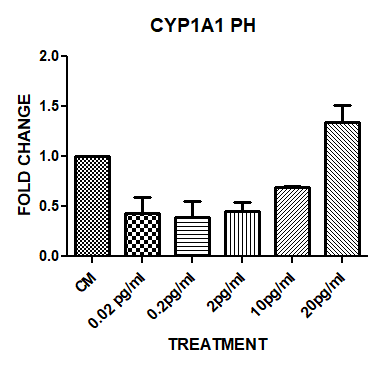

2B

2C

3A

3B

4A

4B

4C

**Figure.5**

**Supplementary figure 3. Fold change graph of *CYP1A1* gene in Hanging drop and polyHEMA culture system for primary buffalo granulosa cells spheroids 3D culture**. The *CYP1A1* expression was upregulated in cultured buffalo granulosa cells than the fresh cells. However, this higher expression was significant (P < 0.05, indicate as ★) only in hanging drop method using exclusively media without containing the isolated milk fat. The expression was found to be non-significant in the remaining all experimental conditions. Nevertheless, the expression of the *CYP1A1* was found to be highest in granulosa cells in both the 3D culture systems, when the cells were treated with the milk fat isolated from the spiked milk containing 0.2pg/ml concentration of TCCD than the cells treated with milk fat without TCDD. 1 and 2 represents experiment results in hanging drop (HD) culture system. 3 and 4 represents experiment results in polyHEMA (PH) culture system. 1 and 3 represent treatment of spheroids with TCDD. 2 and 4 represents treatment of spheroids with fat isolated from TCDD spiked milk. CM represents the cells in media. 0.02pg/ml, 0.2pg/ml, 2pg/ml, 10pg/ml and 20pg/ml indicate the conc. of TCDD. FC represents granulosa cells isolated freshly. MF represents milk fat.

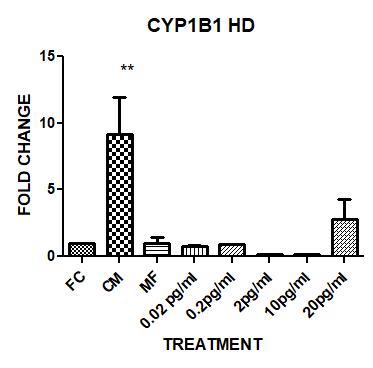

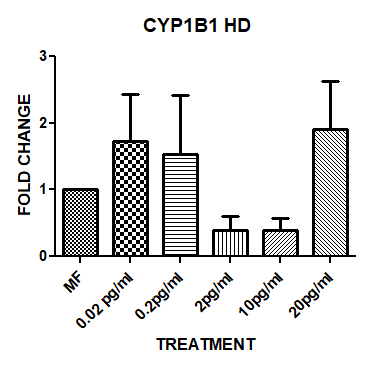

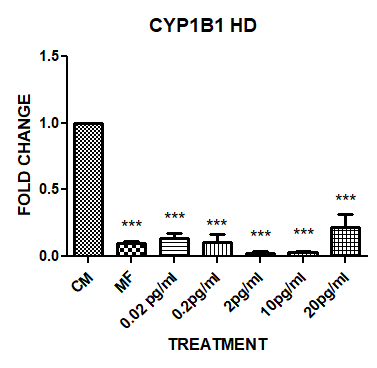

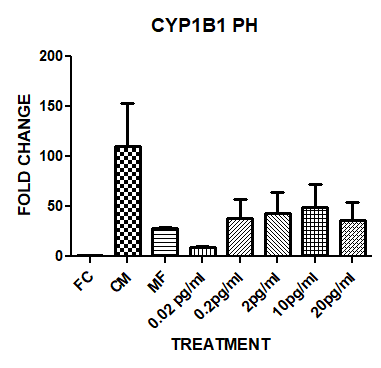

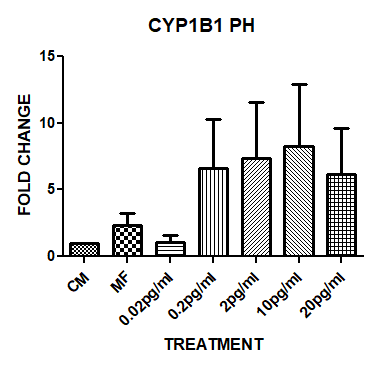

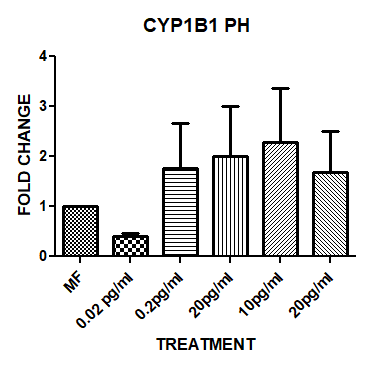

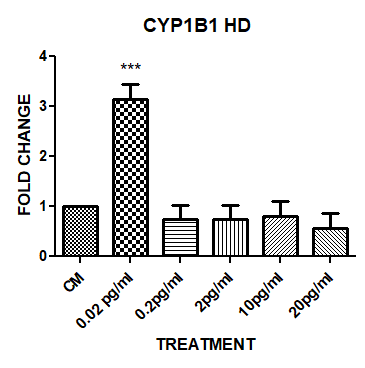

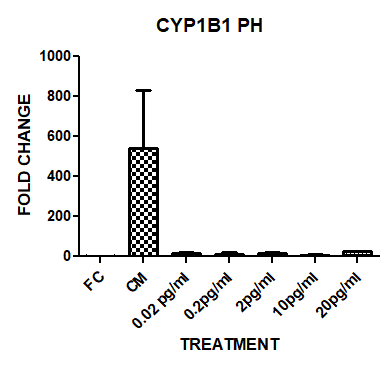

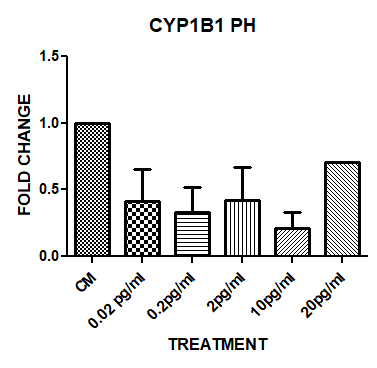

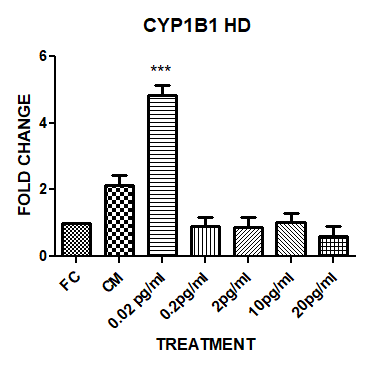

1A

1B

2A

2B

2C

3A

3B

4A

4B

4C

**Supplementary figure 4. Fold change graph of *CYP1B1* gene in Hanging drop and polyHEMA culture system for primary buffalo granulosa cells spheroids 3D culture**. When the buffalo granulosa cell spheroids in hanging drop culture system were treated with media containing directly TCDD, the *CYP1B1* gene expression was significantly (P < 0.05, indicate as ★) increased at 0.02pg/ml concentration than freshly isolated granulosa cells, the cells without TCDD treatment as well as TCDD treatment with other concentrations. However, when the spheroids were treated with the media containing isolated fat from the milk spiked with TCDD, the *CYP1B1* expression was significantly reduced than the cells treated with media itself, but similar to fresh cells. Although there was an increasing trend of *CYP1B1* expression by the fat isolated from the milk spiked with TCDD, there was no significant difference between milk fat treatment and the treatment with milk fat containing TCDD in hanging drop culture system. Though a similar trend was observed in polyHEMA culture system, there was no significant difference between milk fat treatment and the treatments with milk fat containing TCDD. These observations indicate that the cells might have adopted tolerance to TCDD treatment regarding *CYP1B1* expression in 24 hours. 1 and 2 represents experiment results in hanging drop (HD) culture system.3 and 4 represents experiment results in polyHEMA (PH) culture system.1 and 3 represent treatment of spheroids with TCDD. 2 and 4 represents treatment of spheroids with fat isolated from TCDD spiked milk. CM represents the cells in media. 0.02pg/ml, 0.2pg/ml, 2pg/ml, 10pg/ml and 20pg/ml indicate the conc. of TCDD. FC represents granulosa cells isolated freshly. MF represents milk fat.

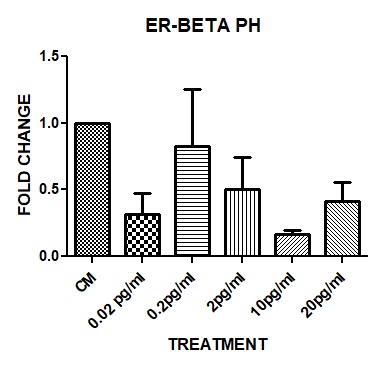

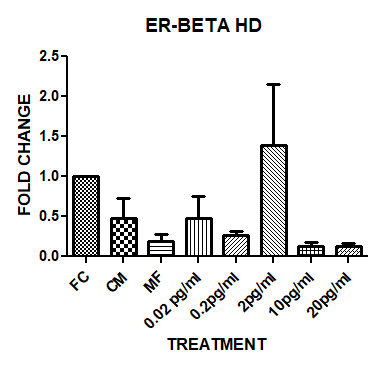

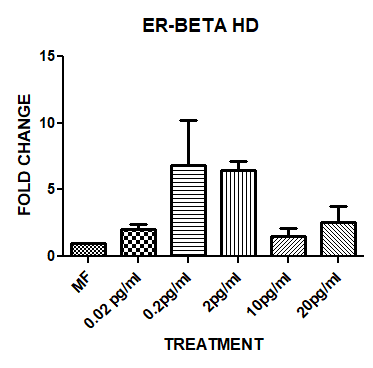

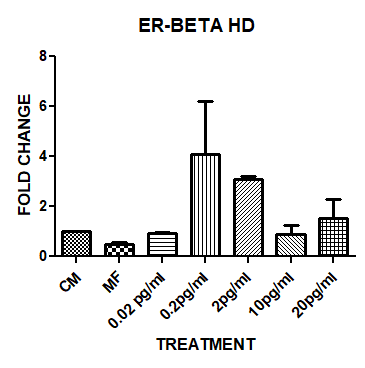

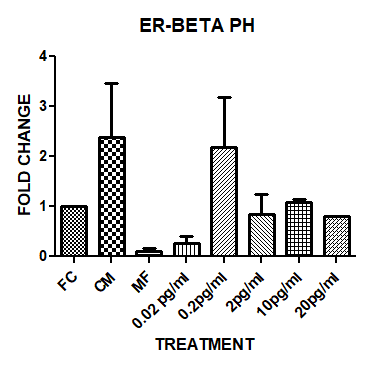

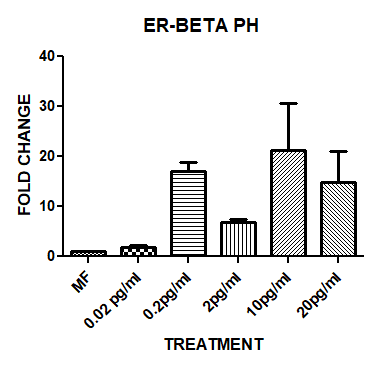

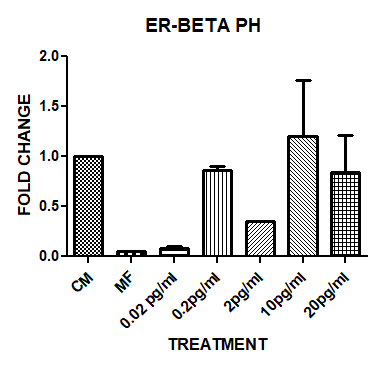

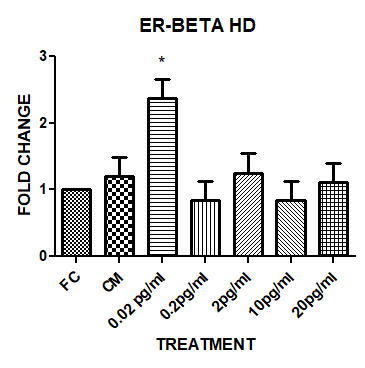

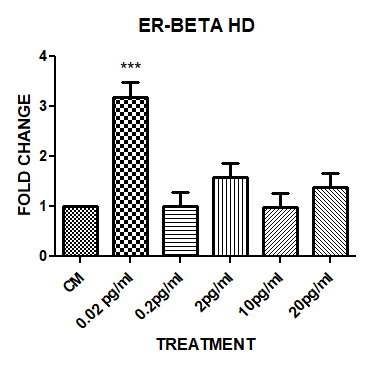

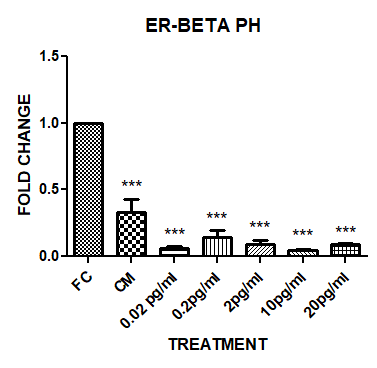

1A

1B

2A

2B

2C

3A

3B

4A

4B

4C

**Supplementary figure 5. Fold change graph of *ER-beta* gene in Hanging drop and polyHEMA culture system for primary buffalo granulosa cells spheroids 3D culture**. When the buffalo granulosa cell spheroids in hanging drop culture system were treated with TCDD directly in the media, the expression of the *ER-β* gene was significantly (P < 0.05, indicate as ★) higher at 0.02pg/ml TCDD concentration than fresh and control cultured spheroids, and the expression declined at later concentrations of TCDD treatment (Figure.9-1A and 2A). However, a significantly declining trend of *ER-β* gene expression was found in the cultured spheroids than the fresh cells in polyHEMA system when they were treated with the media containing TCDD directly (Figure.3A). On the contrary, the *ER-β* gene expression was increased, although not significantly, by the milk fat containing TCDD greater than 0.2 pg/ml than the milk fat treatment in both the culture systems (Figure.9-2B, 2C, 4B and 4C). These observations indicate that the *ER-β* could be another potential gene marker to identify the dioxins in milk. 1 and 2 represents experiment results in hanging drop (HD) culture system. 3 and 4 represents experiment results in polyHEMA (PH) culture system. 1 and 3 represent treatment of spheroids with TCDD. 2 and 4 represents treatment of spheroids with fat isolated from TCDD spiked milk. CM represents the cells in media. 0.02pg/ml, 0.2pg/ml, 2pg/ml, 10pg/ml and 20pg/ml indicate the conc. of TCDD. FC represents granulosa cells isolated freshly. MF represents milk fat.

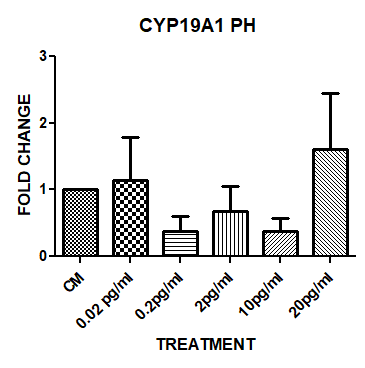

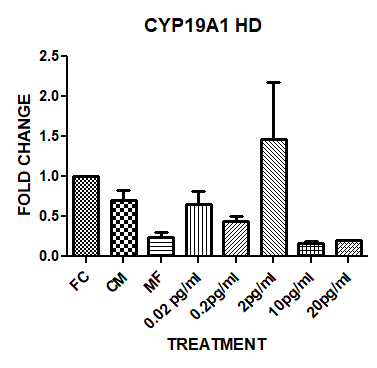

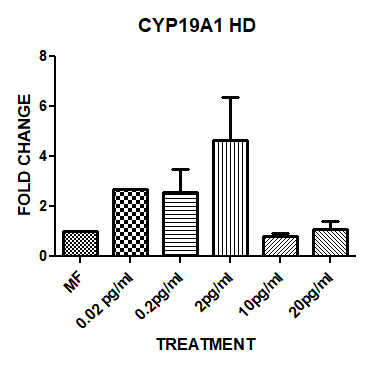

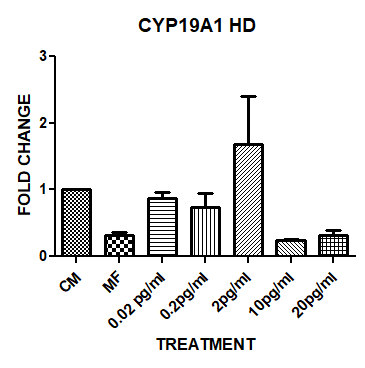

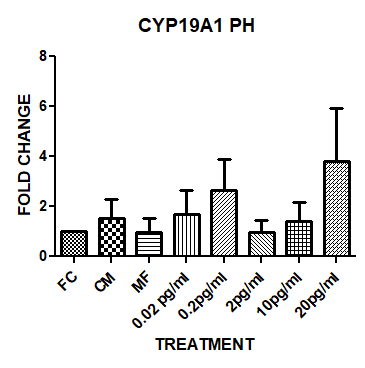

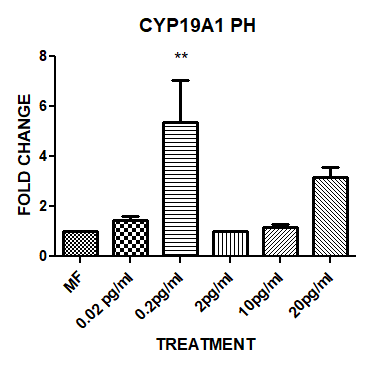

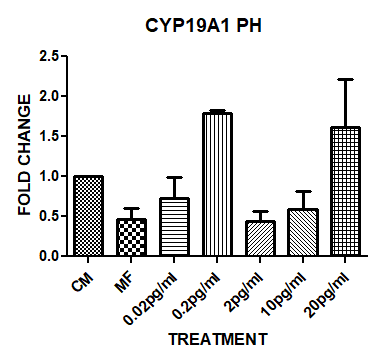

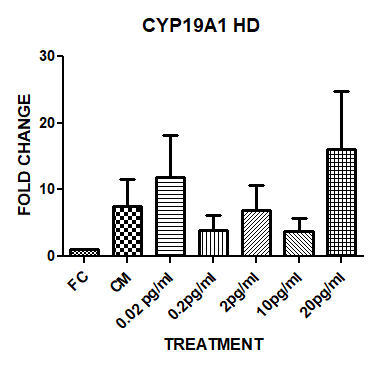

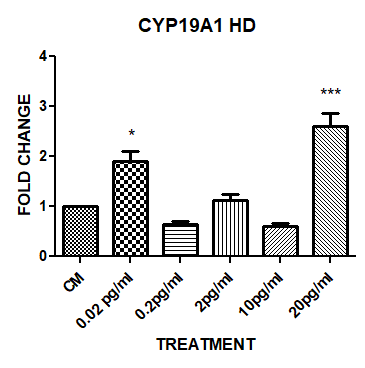

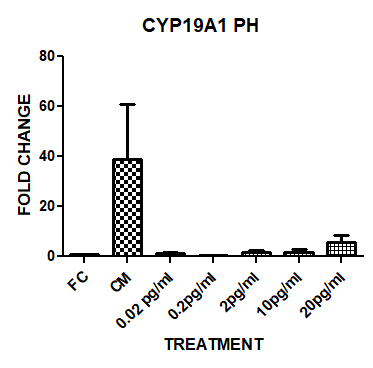

1A

1B

2A

2B

2C

3A

3B

4A

4B

4C

**Supplementary figure 6. Fold change graph of *CYP19A1* gene in Hanging drop and polyHEMA culture system for primary buffalo granulosa cells spheroids 3D culture**. A significant (P < 0.05, indicate as ★) increase of the *CYP19A1* expression was found by 0.02 and 20 pg/ml TCDD concentrations than the cultured cells without TCDD treatment directly in hanging drop culture system. However, when these spheroids in hanging drop culture system were treated with the milk fat containing TCDD, 2pg/ml of TCDD enhanced the *CYP19A1* gene expression though it was not significant than the spheroids treated with only milk fat. In polyHEMA system, when these spheroids were treated with the milk fat containing TCDD, 0.2pg/ml of TCDD enhanced the *CYP19A1* gene expression than those spheroids treated with milk fat alone and the milk fat spiked with other concentrations of TCDD. While in all other conditions there was no significant difference in the *CYP19A1* expression. 1 and 2 represents experiment results in hanging drop (HD) culture system. 3 and 4 represents experiment results in polyHEMA (PH) culture system. 1 and 3 represent treatment of spheroids with TCDD. 2 and 4 represents treatment of spheroids with fat isolated from TCDD spiked milk. CM represents the cells in media. 0.02pg/ml, 0.2pg/ml, 2pg/ml, 10pg/ml and 20pg/ml indicate the conc. of TCDD. FC represents granulosa cells isolated freshly. MF represents milk fat.

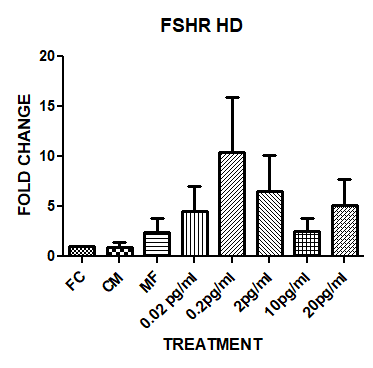

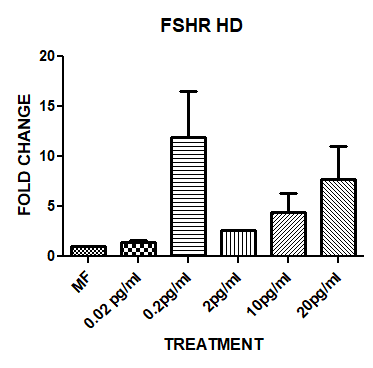

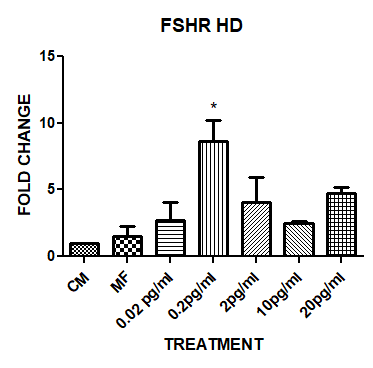

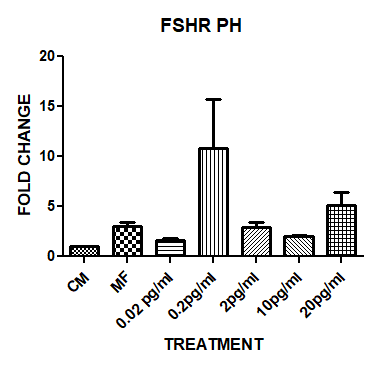

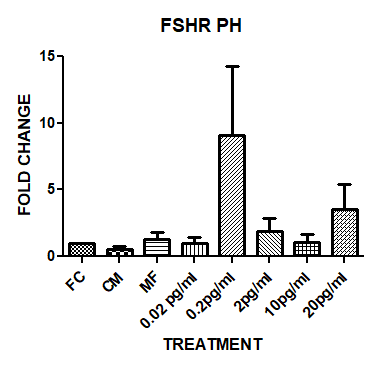

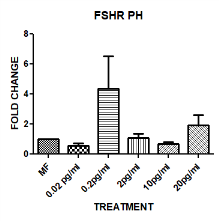

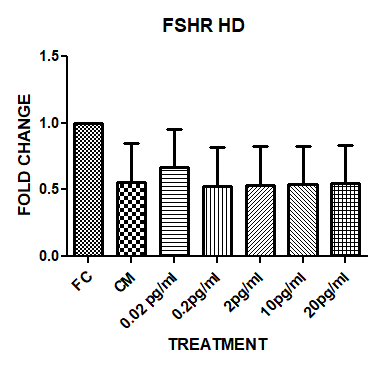

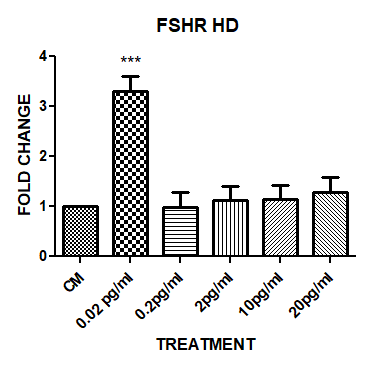

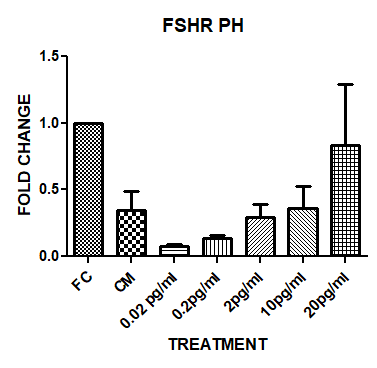

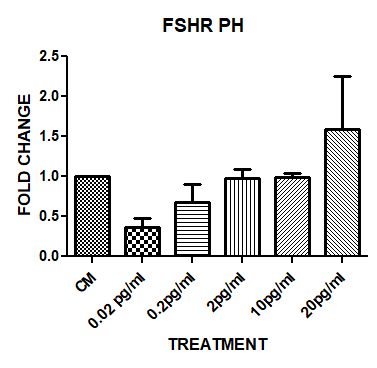

1B

2C

4C

2B

1A

2A

3A

3B

4A

4B

**Supplementary figure 7. Fold change graph of *FSHR* gene in Hanging drop and polyHEMA culture system for primary buffalo granulosa cells spheroids 3D culture**. When the buffalo granulosa cell spheroids in hanging drop culture system were treated with TCDD directly in media, the expression of FSHR gene was still maintained similar to fresh cells (Figure.10-1A). However, its expression was significantly (P < 0.05, indicate as ★) higher at 0.02pg/ml of TCDD than the control cultured spheroids and other concentrations of TCDD in the media (Figure.10-1B). On the contrary, 20 pg/ml of TCDD present directly in the media enhanced, although not significant, the FSHR expression than the cultured control granulosa cell spheroids in polyHEMA culture system (Figure.10-3A and 3B). Interestingly, milk fat containing 0.2 pg/ml of TCDD increased the FSHR expression (not significant) than the treatment of milk fat alone and the milk fat containing other concentration of TCDD in both hanging drop and polyHEMA culture systems. 1 and 2 represents experiment results in hanging drop (HD) culture system. 3 and 4 represents experiment results in polyHEMA (PH) culture system. 1 and 3 represent treatment of spheroids with TCDD. 2 and 4 represents treatment of spheroids with fat isolated from TCDD spiked milk. CM represents the cells in media. 0.02pg/ml, 0.2pg/ml, 2pg/ml, 10pg/ml and 20pg/ml indicates the conc. of TCDD. FC represents granulosa cells isolated freshly. MF represents milk fat

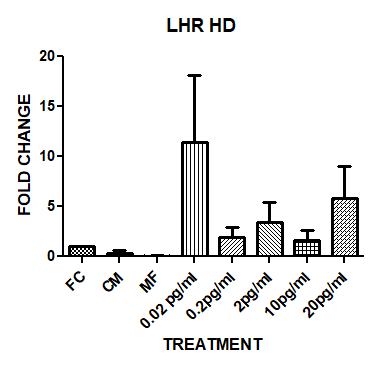

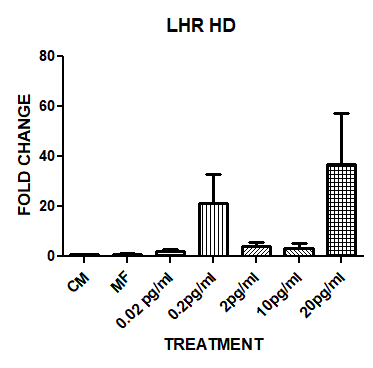

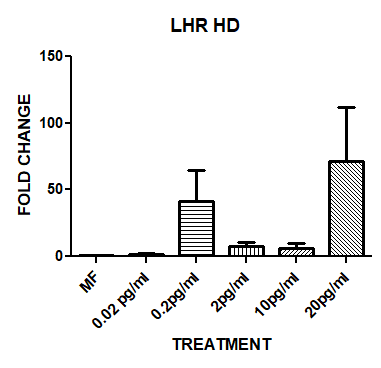

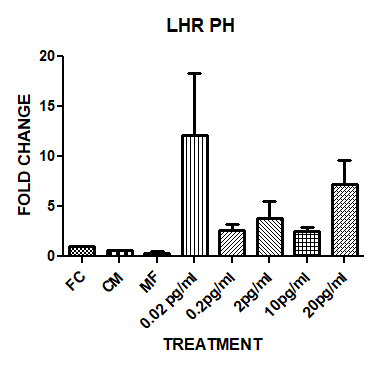

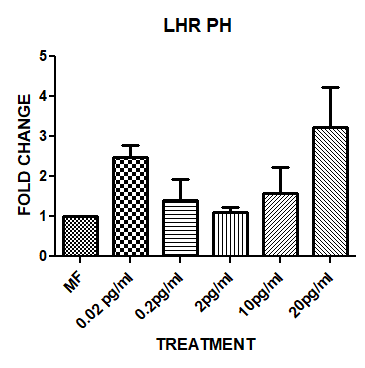

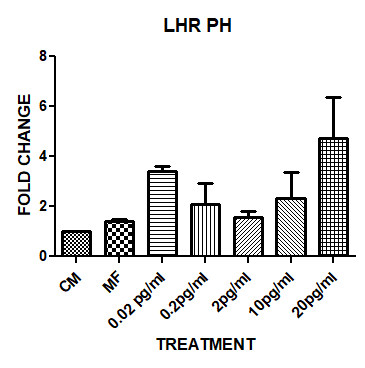

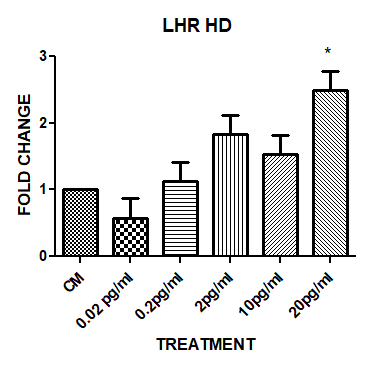

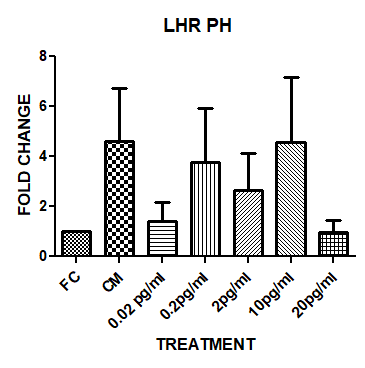

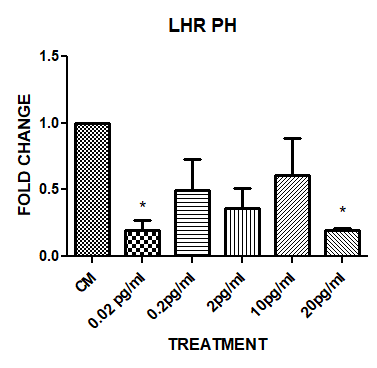

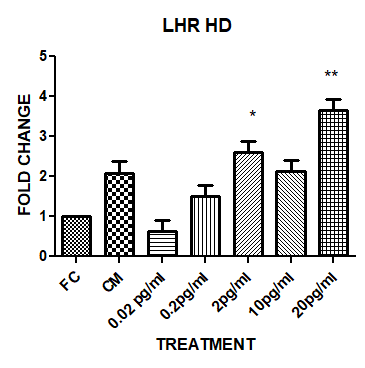

1A

1B

2A

2B

2C

3A

3B

4A

4B

4C

**Supplementary figure 8. Fold change graph of *LHR* gene in Hanging drop and polyHEMA culture system for primary buffalo granulosa cells spheroids 3D culture**. *LHR* gene was found to be gradually upregulated in a dose dependent manner of TCDD in hanging drop culture system when the spheroids of buffalo granulosa cells were treated with the media containing TCDD directly. However, its expression was decreased in polyHEMA culture system in a similar experiment. Although it was not significant (P < 0.05, indicate as ★), the LHR gene expression was higher in those spheroids of buffalo granulosa cells treated with milk fat containing either 0.2pg/ml or 20 pg/ml of TCDD than milk fat treatment alone. A similar trend was also observed in the polyHEMA culture system by the treatment of milk fat containing 0.02pg/ml and 20 pg/ml of TCDD. 1 and 2 represents experiment results in hanging drop (HD) culture system. 3 and 4 represents experiment results in polyHEMA (PH) culture system. 1 and 3 represent treatment of spheroids with TCDD. 2 and 4 represents treatment of spheroids with fat isolated from TCDD spiked milk. CM represents the cells in media. 0.02pg/ml, 0.2pg/ml, 2pg/ml, 10pg/ml and 20pg/ml indicate the conc. of TCDD. FC represents granulosa cells isolated freshly. MF represents milk fat.

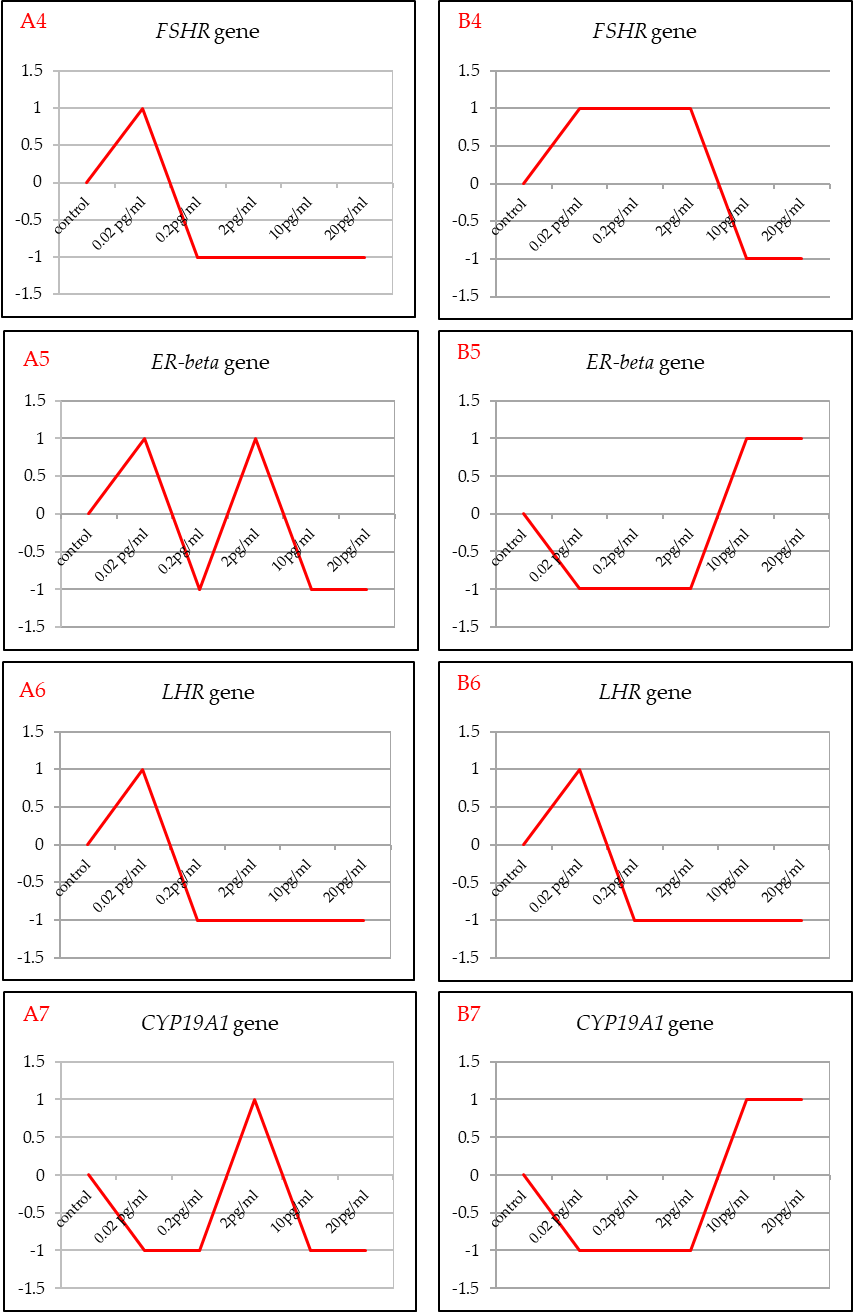

Correlation

Correlation

Correlation

Correlation

PolyHEMA culture system

Hanging drop culture system

**Supplementary figure 9. Correlation analysis for the expression of the selected granulosa cell marker genes’ between with direct TCDD treatment and TCDD treatment through milk fat for the 3D cultured primary buffalo granulosa cells spheroids.** A and B represent the correlation analysis between the above two kinds of treatments in hanging drop (HD) and polyHEMA culture systems, respectively. The No.s 4, 5, 6 and 7 indicate the FSHR, ER-beta, LHR and CYP19A1 gene expressions, respectively. The X-axis shows different concentrations of TCDD and the Y-axis indicates the correlation value between the two kinds of TCDD treatments. The correlation analysis revealed that no perfect positive correlation of granulosa cell specific transcripts between the two kinds of TCDD treatments, indicating the effect of milk fat.
